# Structural characterization and extended substrate scope analysis of two Mg^2+^-dependent O-methyltransferases from bacteria

**DOI:** 10.1101/2023.01.28.526015

**Authors:** Nika Sokolova, Lili Zhang, Sadaf Deravi, Rick Oerlemans, Matthew R. Groves, Kristina Haslinger

## Abstract

Oxygen-directed methylation is a ubiquitous tailoring reaction in natural product pathways catalysed by O-methyltransferases (OMTs). Promiscuous OMT biocatalysts are thus a valuable asset in the toolkit for sustainable synthesis and optimization of known bioactive scaffolds for drug development. Here, we characterized two bacterial OMTs from *Desulforomonas acetoxidans* and *Streptomyces avermitilis* in terms of their enzymatic properties and substrate scope and determined their crystal structures. Both OMTs methylated a wide range of catechol-like substrates, including flavonoids, coumarins, hydroxybenzoic acids and their respective aldehydes, an anthraquinone and an indole. One enzyme also accepted a steroid. The product range included pharmaceutically relevant compounds such as (iso)fraxidin, iso(scopoletin), chrysoeriol, alizarin 1-methyl ether and 2-methoxyestradiol. Interestingly, certain non-catechol flavonoids and hydroxybenzoic acids were also methylated. This study expands the knowledge on substrate preference and structural diversity of bacterial catechol OMTs and paves the way for their use in (combinatorial) pathway engineering.

**Table of contents:** **Two promiscuous O-methyltransferases** from bacteria were found to methylate a panel of catechol substrates towards high-value medicinal compounds. Surprisingly, the non-catechol substrates 5-hydroxyflavonoids and *o*-hydroxybenzoic acids/aldehydes were also methylated at low conversion rates. The crystal structures reveal potential target sites for enzyme engineering for biocatalytic applications.

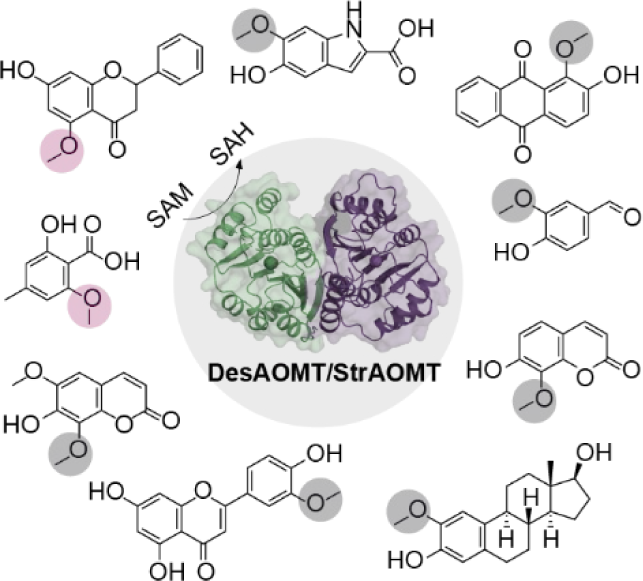

## Introduction

Methylation is a common modification of all major classes of secondary metabolites and alters their physicochemical properties and biological activity. In particular, oxygen-directed methylation (O-methylation) of natural product scaffolds increases their lipophilicity, introduces steric effects that may affect conformation, and stabilizes reactive intermediates in multistep biosynthetic pathways^[1,2]^. Accordingly, this “methyl effect” is widely used in medicinal chemistry to increase membrane permeability, bioavailability and stability of lead compounds in drug development and modulate their target-binding properties^[2,3]^.

In nature, methylation of hydroxyl groups is carried out by O-methyltransferases (OMTs) – a diverse family of enzymes relying primarily on S-adenosylmethionine (SAM) as methyl donor. One subgroup of secondary metabolite methyltransferases known as Class I or catechol OMTs (COMTs) catalyses methylation of phenols bearing vicinal hydroxyl groups (catechols), often with a preference for one of the two positions^[2]^. The resulting mono-O-methylated catechol group, the guaiacol, is a recurring motif in many plant natural products of pharmaceutical and nutraceutical interest, including vanillin, eugenol, capsaicin and methylated flavonoids. COMTs are also well-studied in animals due to their role in the inactivation of catecholamine neurotransmitters and xenobiotics^[4]^. One characteristic of COMTs is their dependence on the binding of a divalent cation, usually Mg^2+^, for full catalytic activity. Metal-independent or Class II OMTs, on the other hand, utilize a catalytic base for deprotonation of the target hydroxyl group and have a broader substrate scope than Class I OMTs^[5]^.

While best studied in plants and animals, class I OMTs are also ubiquitous in bacteria. In some cases, they are encoded in secondary metabolite biosynthetic gene clusters (BGCs), where they methylate precursors of complex antibiotic agents such as the L-DOPA building block in saframycin MX1^[4]^ or 4,5-dihydroxyanthranilic acid in tomaymycin^[7]^. The guaiacol group is also present in limazepines^[8]^ and streptonigrin^[9]^, although the OMTs from the corresponding BGCs have not been characterized. The majority of known bacterial COMTs, however, are encoded outside of BGCs, and their cellular targets and physiological functions remain elusive. Several *in vitro* activity studies demonstrate a high tolerance of bacterial COMTs towards non-natural catechol substrates such as catecholamines, phenylpropanoids and flavonoids^[10–13]^.

Heterologously expressed OMTs have already been successfully integrated in the engineered pathways towards high-value compounds like ferulic acid^[14]^, curcuminoids^[15]^ and vanillin^[16–18]^, with the latter resulting in the establishment of a commercial process. Additionally, a number of recent studies have focused on the fine-tuning of COMT regioselectivity^[12,19]^, but the substrate scope is mostly confined to plant phenylpropanoids and flavonoids. As O-methylation is one of the most common tailoring reactions in natural product biosynthesis alongside hydroxylation, glycosylation and prenylation, promiscuous OMTs are a valuable asset in the toolkit for pathway engineering and diversity-oriented combinatorial biosynthesis^[20]^. To that end, thorough characterization of the substrate and product scope of candidate enzymes is essential to fully exploit their catalytic potential.

Here, we report heterologous expression and *in vitro* characterization of two promiscuous O-methyltransferases from bacteria. We evaluated the biosynthetic potential of these OMTs on a set of representative natural product scaffolds, demonstrating the successful methylation of several non-canonical substrates and revealing the substrate-dependent nature of OMT regioselectivity. The product scope of the OMTs included several compounds of pharmaceutical and nutraceutical relevance. We furthermore determined high-resolution X-ray crystal structures of the OMTs to reveal potential target sites for tuning regioselectivity or enhancing the catalytic efficiency by enzyme engineering.

## Results and Discussion

### Biochemical characterization and *in vitro* activity assays

In a previous study^[14]^, we used cell-free transcription/translation followed by *in vitro* activity testing to screen a panel of putative O-methyltransferases against several catechol-like compounds. Some of these enzymes catalysed regiospecific methylation of caffeic acid to ferulic acid *in vitro* and in an *Escherichia coli* cell factory. From this set of enzymes, we selected two OMTs for further characterization and substrate scope analysis in this study: the top-performing enzyme StrAOMT from *Streptomyces avermitilis* (UniProt accession: Q82B68), which was previously shown to methylate several flavonoids *in vitro*^[10]^, and DesAOMT from *Desulfuromonas acetoxidans* (UniProt accession: Q1JXV1) - an otherwise uncharacterized enzyme. DesAOMT struck us as interesting because of its lower molecular weight and the lack of a putative catalytic triad conserved in all known Class I plant and bacterial OMTs^[21]^. Both proteins belong to the PF01596 family, characterized members of which include mammalian COMTs, plant caffeoyl-CoA and flavonoid OMTs, and several secondary metabolite OMTs from bacteria and fungi.

First, we performed an analysis of the genome neighbourhood^[22]^ of the OMTs and their sequence homologs to check for clues on the natural substrates and functions of these enzymes (Figure S1). We saw that the StrAOMT gene is surrounded, among others, by domains encoding putative lipase, acetyl-CoA acetyltransferase and cholest-4-en-3-one 26-monooxygenase functionalities, which might be indicative of a steroid metabolic pathway, as well as several transporters and a prenyltransferase-like repeat protein in the extended neighbourhood. This motivated us to include a terpene and a steroid derivative in the substrate scope analysis. In a broader phylogenomic analysis, we noticed that the StrAOMT gene neighbourhood is highly conserved in other Streptomyces genomes. The immediate neighbourhood of the DesAOMT gene contains diguanylate cyclase and diguanylate phosphodiesterase domains, which are responsible for the synthesis and degradation of a bacterial messenger cyclic di-GMP (Figure S1). We did not find any literature precedent for an association of such genes with methyltransferases, and this combination of genes does not appear to be conserved among sequence homologues of DesAOMT.

Second, we characterized both enzymes biochemically with caffeic acid as a substrate. We cloned the two genes into pET-21b(+) with a C-terminal 6xHis-tag for expression in *E. coli*. We purified the proteins to homogeneity with a two-step protocol comprising nickel-affinity and size-exclusion chromatography (Fig. S2). Next, we confirmed that the purified enzymes were capable of methylating caffeic acid under the previously used reaction conditions^[14,23]^ to the *meta*- and the *para*-methoxy products, ferulic acid (FA) and iso-ferulic acid (IFA), respectively (Figure 1a). We then set out to investigate their biochemical properties and optimize the reaction conditions. Both enzymes were tolerant to higher reaction temperatures with the maximum catalytic activity at 40°C and 45°C for DesAOMT and StrAOMT, respectively (Figure 2b). We confirmed that both enzymes are dependent on Mg^2+^ for catalytic activity (Figure 2c, d): the addition of Ca^2+^ or ethylenediaminetetraacetic acid (EDTA) fully inhibited both enzymes, whereas we observed weak residual activity in reactions with no additives (“none”). This may be attributed to the presence of residual Mg^2+^ from protein purification and storage. Alkaline conditions were preferred by both enzymes with an optimum at pH 8–8.5 in Tris-HCl buffer (Figure 1e, f). It is noteworthy that increasing the pH prompted a noticeable shift in regioselectivity of StrAOMT, while overall maintaining a preference for the *meta*-isomer (ferulic acid). With a drop in overall activity, StrAOMT appears to become less regiospecific at pH higher than 8.0. DesAOMT showed overall lower regioselectivity that remained stable across a wide pH range. Overall, with the optimized temperature and buffer conditions (37°C, Tris-HCl pH 8) and by using a 5-fold excess of SAM, we achieved ∼85% conversion of caffeic acid by both enzymes in one hour. Higher conversion rates could be achieved by further increasing the concentration of SAM.

**Figure 1.**
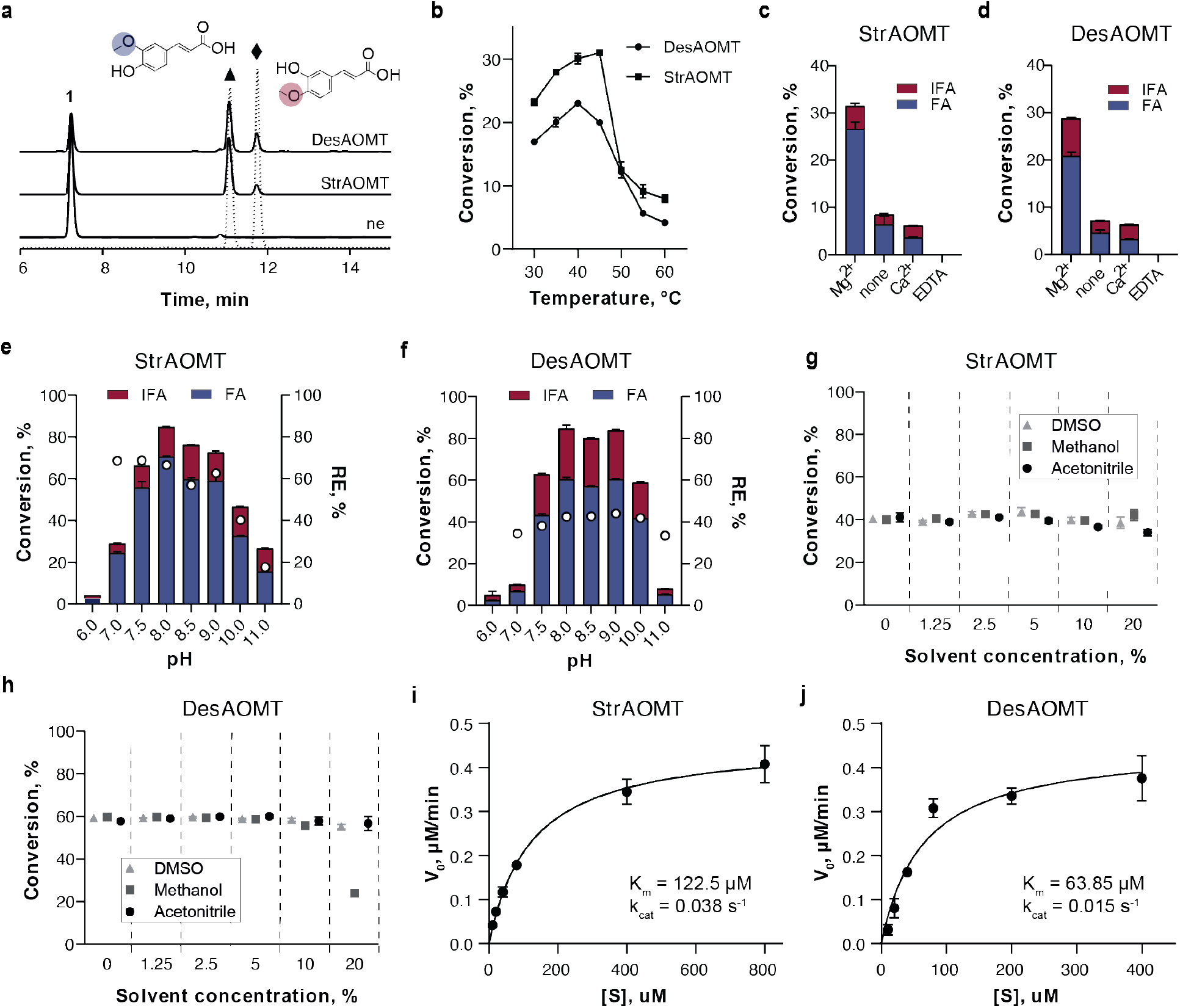
Biochemical properties of DesAOMT and StrAOMT with caffeic acid as substrate. Standard reaction conditions (if not stated otherwise): 20 mM Tris-HCl pH 7.5, 20 mM MgCl_2_, 100 µM caffeic acid, 200 µM SAM, 1 µM enzyme; 1 h at 37°C without shaking. a) Chromatogram of the OMT-catalysed reactions compared to the no enzyme control (“ne”) (λ=310nm); b) substrate conversions at different reaction temperatures; c-d) substrate conversions with added Mg^2+^, Ca^2+^, EDTA, or no additives (“none”); e-f) substrate conversions at different buffer pH (buffers: NaPi pH 6; HEPES pH 7; Tris-HCl pH 7.5, 8, 8.5; Gly-NaOH pH 9, 10, 11); g-h) substrate conversions in the presence of organic solvents; i-j) Michaelis-Menten kinetics of StrAOMT and DesAOMT. The data are represented as mean ± standard deviation of three technical replicates. The full statistical report non-linear regression is shown in Table S1.

**Figure 2.**
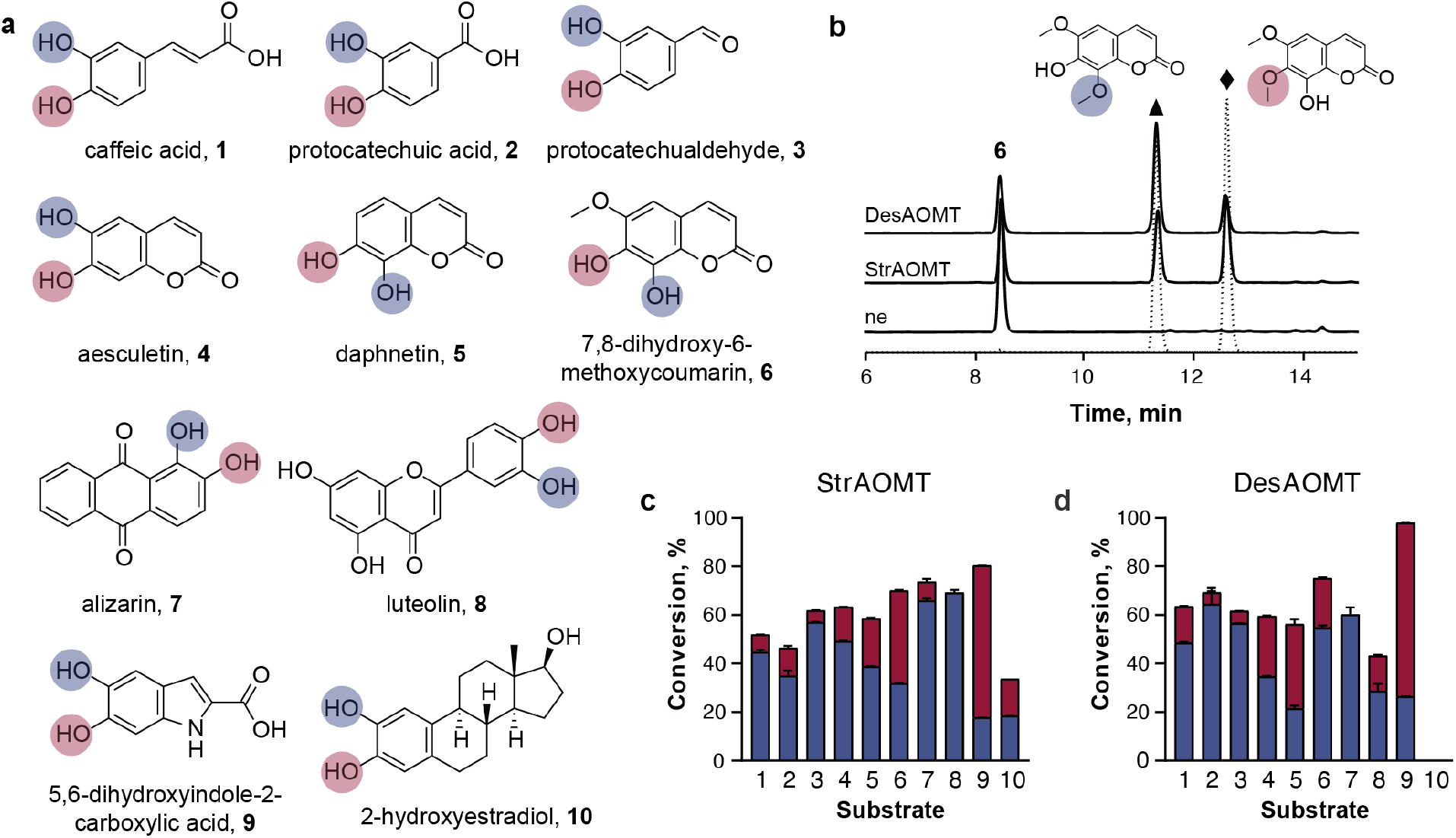
*In vitro* activity and regioselectivity of DesAOMT and StrAOMT with catechol-like compounds. a) Panel of catechol-like substrates highlighting possible O-methylation sites in red and blue; b) HPLC-based identification of reaction products of StrAOMT and DesAOMT exemplified by substrate **6**. Solid lines – reaction products, dashed lines – authentic standards of the potential products isofraxidin (triangle) and fraxidin (diamond), ne – “no enzyme” control; detection at λ = 310 nm; c) and d) conversion of substrates **1-10** into the two possible products depicted as stacked histograms (colour coding according to panel a).

Since several compounds in our intended substrate panel are poorly soluble in water, we sought to explore how tolerant DesAOMT and StrAOMT are to organic solvents. To our surprise, increasing concentrations of dimethylsulfoxide (DMSO) and acetonitrile (ACN) to 20% (v/v) virtually did not affect catalytic activity of the OMTs, while methanol inhibited DesAOMT only at the highest concentration (Figure 1g, h).

Lastly, we determined the apparent Michaelis-Menten kinetic parameters at a fixed SAM concentration of 1 mM at 37°C and pH 7.5. We stopped the reactions after 5, 10, and 15 min and determined the product concentrations by HPLC to estimate the initial reaction velocities using linear regression (Figure S3). Overall, DesAOMT had a lower apparent K_m_ for caffeic acid than StrAOMT, while k_cat_/K_m_ values were similar for the two enzymes (Figure 1i, j). During this series of experiments, we also noticed strong enzyme inhibition at higher substrate concentrations for both enzymes (8000-fold molar excess of substrate over enzyme). A similar observation was previously reported for the bacterial OMT SafC with caffeic acid and dopamine as substrates^[6]^, but the mechanism of this inhibition is not fully understood.

### DesAOMT and StrAOMT methylate a variety of catechol-like scaffolds with differing regioselectivity

Next, we set out to assess the performance of DesAOMT and StrAOMT on a range of catechol-like substrates representative of natural product scaffolds, including flavonoids, coumarins, benzoic and resorcylic acids and their respective aldehydes, an anthraquinone, an indole, a terpene and a steroid (Figure 2a, Figure S4).

We incubated 0.5 mM of the respective substrate with 5 µM DesAOMT or StrAOMT for 16 h at 37°C. We performed the assays at pH 7.5 in view of instability of some substrates in alkaline conditions and added 10-20% DMSO for better solubility of substrates and their methylated products.

Both OMTs exhibited remarkable tolerance towards diverse catechol-like substrates with high conversion rates for substrates **1**-**10** (Figure 2c, d) and moderate to low conversion rates for substrates **17-24** (Figure S4). For the latter substrates, we detected putative methylated products by LC-MS, however, we did not characterize the products any further because the reference compounds were unavailable and the low turnover yields did not warrant in depth structural characterization. Nevertheless, the low level of enzymatic activity observed for these substrates might be an interesting starting point for further investigation.

For substrates **1**-**10** we confirmed the identity of the products by comparing the HPLC retention times and mass over charge values to those of authentic standards (Figure S5) and thereby assessed the regioselectivity of the enzymes. We found that it depends on the chemical scaffold and differs between the two enzymes. Both were selective for the *meta* hydroxyl of phenolic acids and aldehydes **1-3** and aesculetin **4**, but exhibited notable differences when challenged with bulky or highly asymmetric substrates, most notably the coumarins **5** and **6**. StrAOMT was selective for the 8-OH of **5** but methylated its 6-methoxy derivative **6** to a mixture of products with only slight preference for the 7-OH to form fraxidin. On the contrary, DesAOMT was selective for the 7-OH of **5** but produced mostly the 8-O-methylated isofraxidin when challenged with **6** (Figure 2b). Curiously, we observed exclusive conversion of **7** to 3-O-methylated chrysoeriol by both enzymes, which may be attributed to the rigidity and asymmetricity of the molecule. Lastly, StrAOMT also converted the steroid **10** to a mixture of 2- and 3-methoxy products. This is consistent with our analysis of the gene neighbourhood of the StrAOMT encoding gene and may indicate that the enzyme has a natural function in steroid metabolism. Overall, our substrate scope analysis with catechol substrates demonstrates the substrate-dependent nature of COMT regioselectivity *in vitro* and highlights the importance of such studies.

### DesAOMT and StrAOMT also accept non-catechol substrates

After we characterized the substrate scope of both enzymes for the typical catechol-like substrates, we turned to non-canonical phenolic substrates (Figure S4, **26**-**31**). As expected, many compounds were not methylated, such as those with a single (m-coumaric acid) or several *meta*- or *para*-positioned hydroxyl groups (resorcinol, hydroquinone), and phenols with other vicinal substitutions (4-chloro-3-hydroxybenzoic acid, 3-hydroxy-4-methylbenzoic acid). This is probably due to their inability to adopt the proper orientation in the active site and/or coordinate the Mg^2+^ effectively. Similarly, isoferulic acid was not further converted into the dimethylated product.

Unexpectedly, our LC-MS results suggest that StrAOMT and DesAOMT also accept non-catechol flavonoids **11**-**13** (chrysin, pinocembrin and naringenin), albeit with low (<10%) conversion rates. Additionally, StrAOMT can methylate **14-16** (p-orsellinic acid, orcinaldehyde and 2,6-dihydroxybenzoic acid) with low conversion rates (Figure 3a).

**Figure 3.**
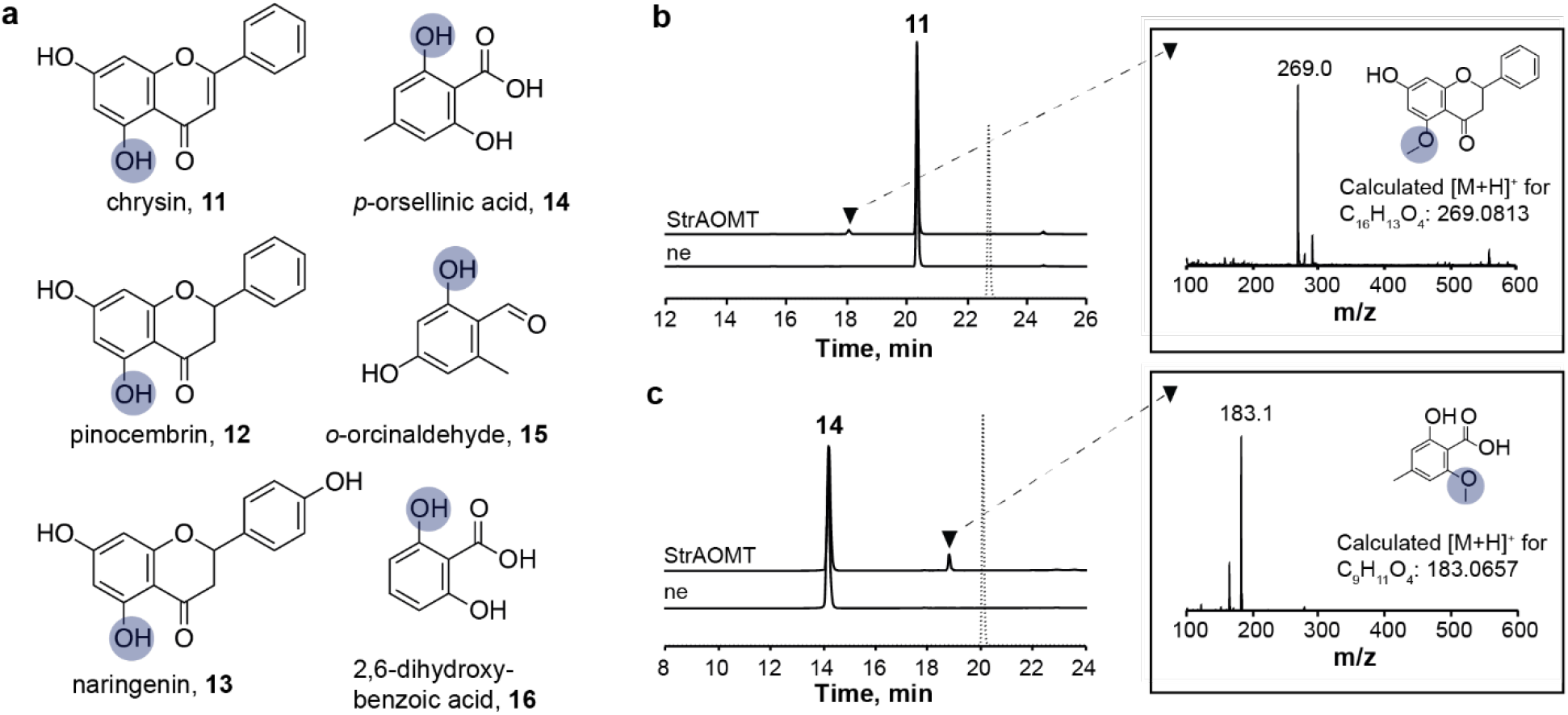
Non-catechol substrates accepted by StrAOMT (**11**-**16**) and DesAOMT (**11**-**13**). a) Panel of non-catechol substrates highlighting putative O-methylation sites in blue; b) and c) elution profiles of StrAOMT-catalysed reactions of **11** and **14** compared to “ne” control and reference compounds of the opposite regioisomers (dashed line); inserts: mass spectra of the putative product peaks (triangle). Detection was performed at λ = 310 nm. Dimethylated products were not observed.

The methylated products of flavonoids **11**-**13** eluted earlier than their substrates (Figure 3b and S6), which is atypical since methylation generally increases hydrophobicity of a given compound. Such an elution profile is consistent with literature reports of 5-O-methylated flavonoids^[24]^, suggesting that **11**-**13** are methylated to chrysin-5-methylether, alpinetin and naringenin-5-methylether, respectively. This assumption is further supported by the fact that the techtochrysin standard – the only other possible methylation product of **11** – does not coelute with the reaction product peak (Figure 3b). Similarly, the product of **14** does not coelute with the readily available authentic standard of the corresponding methyl ester (Figure 3c), which suggests that **14** is methylated at either of the two phenolic hydroxyl groups. Based on the inability of DesAOMT and StrAOMT to accept **25-27**, it is plausible that **15** and **16** are also methylated at the *o*-hydroxyl position.

Thus far, methylation of *o*-hydroxybenzoic acids in natural products has only been described for metal-independent enzymes following a different mechanism, as exemplified by calicheamicin orsellinate 2-O-methyltransferase^[25]^. The biosynthesis of 5-methoxyflavonoids remains elusive. Therefore, our findings for these two bacterial COMTs are rather unusual. Mechanistically, however, it is possible that some non-catechol substrates can chelate Mg^2+^ and thus serve as substrates for COMTs, as exemplified by the potent human COMT inhibitors hydroxyquinoline and tropolone^[4]^. In addition, the observed 5-O-methylation of flavonoids may be attributed to enolization of the C-4 carbonyl in a conjugated flav(an)one system, with the resulting hydroxyl group participating in Mg^2+^ coordination.

In order to elucidate the basis of the differing substrate scope and regioselectivities of DesAOMT and StrAOMT, we next turned to the structural characterization of these enzymes.

### Crystal structures of StrAOMT and DesAOMT

We determined the crystal structures of apo-DesAOMT at 1.5 Å, apo-StrAOMT at 1.5 Å and SAH-bound StrAOMT at 1.8 Å resolution (Table S2). One asymmetric unit (ASU) of the ligand-free StrAOMT contains two copies of the dimer forming an interface in the substrate-binding site, while a canonical COMT dimer is present in the ASU of the SAH-bound structure (Figure 4a). The crystals of the latter were obtained through co-crystallization of StrAOMT with SAM, Mg^2+^ and caffeic acid, suggesting that the enzymatic reaction proceeded *in situ* and the co-product, SAH, remained bound in the active site. Each StrAOMT monomer in both structures adopts the Rossmann fold characteristic of SAM-binding proteins, with seven core β-strands surrounded by eight α-helices. The conformation of the apo- and the ligand-bound form of StrAOMT is highly similar, with an average Cα RMSD of 0.451 between the monomers. Most notably, SAH binding induces conformational changes in the loop region between α2 and α3 adjacent to SAH (Figure 4b). A search^[26]^ for structural homologs of the ligand-free structure of StrAOMT identified *Bacillus cereus* BcOMT2 as the top hit (PDB: 3DUW, Z-score 36.6, RMSD 1Å), closely followed by NkCOMT from *Niastella koreensis* (PDB: 7CVX), TomG from *Streptomyces regensis* (PDB: 5N5D), a putative OMT from *Klebsiella pneumoniae* (PDB: 3TWF) and Rv0187 from *Mycobacterium tuberculosis* (PDB: 6JCL) (Table S3). The largest conformational diversity between these structures is observed in the so-called insertion loop between β5 and α8 (Figure 4d) – a region implicated in binding Coenzyme A and substrate specificity in plant COMTs.

**Figure 4.**
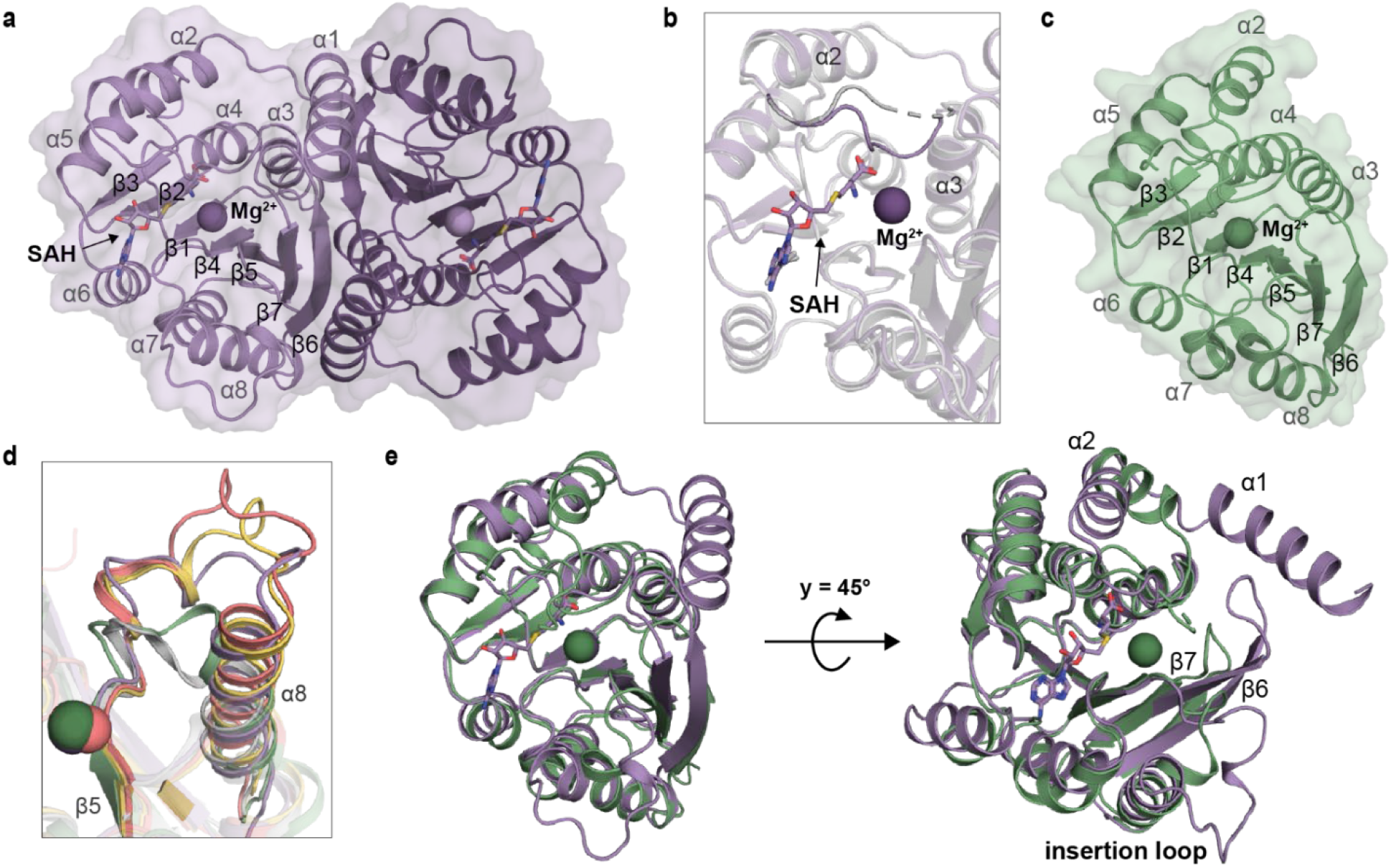
Crystal structures of StrAOMT and DesAOMT. Cartoon representation of a) SAH-bound StrAOMT dimer (holo-, PDB: 8C9S); b) superimposed structures of apo- (grey, PDB: 8C9T) and holo-StrAOMT (purple)zoomed in on the α2-α3 loop adjacent to the ligands ; c) apo-DesAOMT monomer (PDB: 8C9V); d) comparison of the insertion loops (β5-α8) across DesAOMT (green), StrAOMT (purple), LiOMT (yellow, PDB: 2HNK), SafC (pink, PDB: 5LOG) and rat COMT (grey, PDB: 1H1D); e) superimposed monomer structures of holo-StrAOMT (purple) and DesAOMT (green).

The ASU of DesAOMT contains a single monomer (Figure 4c), but the canonical dimer interface can be identified between two adjacent ASUs. Furthermore, we believe that DesAOMT forms a dimer in solution based on its elution profile during size exclusion chromatography (Figure S2). The otherwise canonical Rossman fold of DesAOMT is devoid of the N-terminal α-helix, which is present in all known COMT structures determined to date and has been implicated in catalytic activity^[27]^ and dimerization^[28]^. In general, DesAOMT shares little structural similarity with characterized OMTs, the top hit being LiOMT from *Leptospira interrogans* (PDB: 2HNK, Z-score 26.1, RMSD 1.8 Å), followed by putative OMTs from *Coxiella burnetii* (PDB: 3TR6, Z-score 25.2, RMSD 1.8 Å) and *Mycobacterium tuberculosis* (PDB: 5×7F, Z-score 25.1, RMSD 2 Å), SafC from *Myxococcus xanthus* (PDB: 5LOG, Z-score 25.1, RMSD 1.9 Å) and Rv0187, which is also a structural homolog of StrAOMT. Apart from the missing N-terminal helix, a distinct structural feature of DesAOMT is a short insertion loop typical of animal rather than bacterial COMTs (Figure 4d). When comparing with the StrAOMT structures, the RMSD of Cα is even higher (2.2 Å) and several conformational differences are apparent (Figure 4e). The most noticeable ones are in the insertion loop and the β6-β7 loop regions.

### Active site architecture of StrAOMT and DesAOMT

Both DesAOMT and StrAOMT possess a metal-binding site conserved across all COMTs: D129, D155, N156 in the former and D141, D167, N168 in the latter (Figure S9). In the DesAOMT and holo-StrAOMT structures, we observed electron density in the expected position between these conserved residues and interpreted it as Mg^2+^. An interpretation as Ca^2+^ would also be possible for the DesAOMT structure, since this enzyme was crystallized in a buffer with a mix of divalent metal ions. There might in fact be a mix of ions occupying the binding sites throughout the crystal of DesAOMT. In the holo-StrAOMT structure, we observed additional electron density that we interpreted as 2-pyrrolidone that likely originated from the crystallization solution. The position of this electron density is not near the catalytic residues and is unlikely to be representative of a substrate- or product-bound state.

Despite numerous attempts, co-crystallization and soaking of StrAOMT with caffeic acid or aesculetin failed to yield substrate-bound crystals. Therefore, we compared the holo-StrAOMT structure to that of its structural homolog NkCOMT, which is complexed with the substrate protocatechuic acid (PDB: 7CVX, chain A) (Figure 5a). The positions of Mg^2+^ and SAH are almost identical in holo-StrAOMT and NkCOMT, as are the residues lining the active sites of the two enzymes. Apart from the Mg^2+^ ion, catalytic activity of known catechol OMTs seems to rely on the absolutely conserved active site triad K-N-D, which is thought to facilitate deprotonation of the aromatic substrate prior to methyl transfer^[21]^. K144, N168, and D215 of StrAOMT are aligned with K142, N166, and D213 of NkCOMT, respectively, forming a typical catalytic triad in the active site. Additionally, K212 of StrAOMT is aligned with K210 of NkCOMT, which closely approaches the 4-hydroxyl group of the substrate. This double-lysine arrangement is shared by other structural homologs of StrAOMT (PDB: 3DUW, 6JCL and 3CBG) (Figure S9). The importance of this conservation is stressed by the fact that the 4-hydroxyl binding lysine of 3CBG, which was also shown to be essential for catalytic activity^[27]^, is residue three of the amino acid chain and is brought into the active site by the N-terminal loop. The aromatic ring of dihydroxybenzoic acid in NkCOMT is sandwiched between I39 and R169, which are aligned with I41 and R171 in StrAOMT.

**Figure 5.**
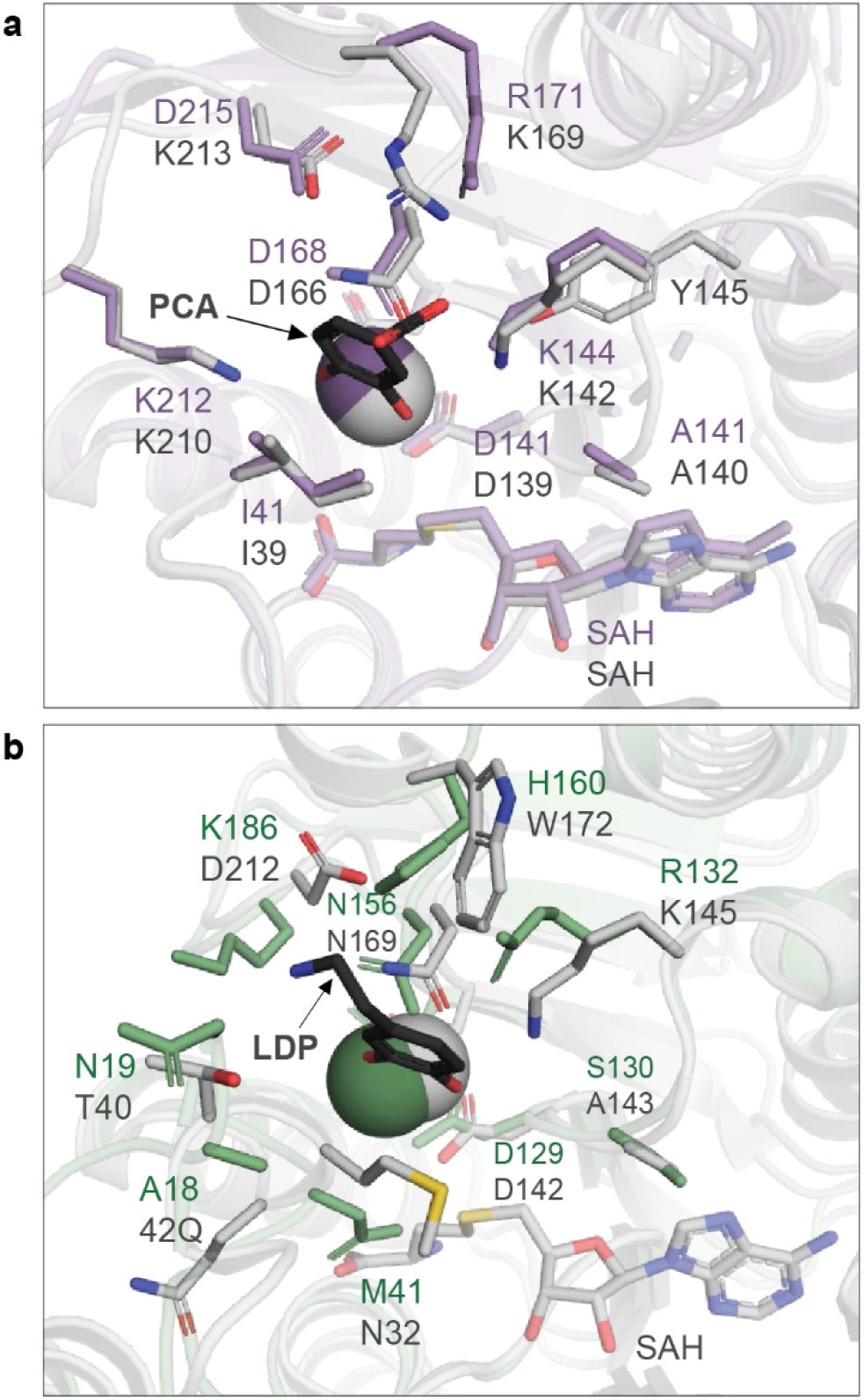
Active site architectures and substrate-binding pockets of DesAOMT and StrAOMT. a) Superimposed structures of holo-StrAOMT (purple) and NkCOMT (grey) complexed with protocatechuic acid (PCA) (PDB: 7CVX); b) superimposed structures of DesAOMT (green) and SafC (grey) complexed with dopamine (LDP) (PDB: 5LOG).

The closest substrate-bound structural homolog of DesAOMT is SafC complexed with dopamine (PDB: 5LOG). It must be noted that the unresolved portion of the loop between α2 and α3 imposes limitations on the examination of the active site of DesAOMT; however, several distinct structural features are apparent. As predicted, R132 of DesAOMT is aligned with K145 of SafC, which was shown to be essential for its catalytic activity, while the remaining putative catalytic residues N169 and D212 are aligned with N156 and K186 in DesAOMT. The latter is located within 5 Å of the putative substrate position and might interact with its other hydroxyl group provided that SAM-induced conformational changes bring it closer to Mg^2+^ (as observed for SAH-bound StrAOMT). DesAOMT is thus the first characterized COMT bearing an arginine in place of the absolutely conserved active site lysine. In the enzyme similarity network generated by Haslinger et al. (Figure S7), the subcluster harbouring DesAOMT (mostly featuring sequences from extremophile bacteria) shares its characteristic features: the active site R-N-K triad, a short insertion loop and a missing N-terminal α-helix (Figure S8). Taken together, these observations might be hinting at a new, possibly more ancient, subgroup of bacterial COMTs.

While in plant COMTs the conserved catalytic triad K-N-D was deemed essential for catalysis^[21]^, there is no clear consensus on the involvement of the catalytic residues in bacterial COMTs. It is generally assumed that the conserved lysine facilitates deprotonation of a hydroxyl group of the substrate, but results of mutagenesis studies are often inconsistent, ranging from a complete loss of activity^[23]^ to its slight reduction with a change in the regioselectivity profile^[11]^. In SynOMT, on the other hand, mutating K3, which is not part of the assumed catalytic triad and is structurally conserved only in a subgroup of bacterial COMTs, completely abolished activity. The unique active site architecture of DesAOMT adds to the long-standing ambiguity surrounding the reaction mechanism of bacterial COMTs, calling for dedicated mechanistic studies like the ones conducted for plant COMTs.

### Structural determinants of substrate specificity and regioselectivity

In model COMT structures, the catalytic Mg^2+^ is located at the bottom of a deep groove lined predominantly with hydrophobic residues, which a catechol substrate can penetrate in two possible orientations. Catechols with polar or ionizable side chains are more likely to orient towards the solvent, while substrates with more hydrophobic substituents may favour orientation towards the “hydrophobic wall” of the enzyme; this results in *meta*- or *para*-selective methylation, respectively^[29]^. Thus, the regioselectivity of COMT-catalysed methylation is largely dictated by the chemical nature of the substrate, which is also apparent from the regioselectivity patterns of DesAOMT and StrAOMT. For instance, **6** differs from **5** only by an 8-methoxy group, yet StrAOMT exhibits opposite regioselectivities with these two substrates.

Accordingly, engineering efforts to modulate the regioselectivity of COMTs have focused mainly on mutating the residues lining the catechol-binding pocket. A notable example is the Y51R mutation in PFOMT, which alone led to the production of a 1:1 mixture of methylated eriodyctiol products (as opposed to the exclusive *meta* methylation of the wild-type enzyme), while complete transition to *para* selectivity was achieved by an additional N202W mutation at the opposite site of the catechol pocket^[19]^. In that light, some of the promising candidates for site-directed mutagenesis include R171, I41 and D215 in StrAOMT or H160, A18 and K186 in DesAOMT.

Interestingly, despite having highly similar active site architectures, NkCOMT and StrAOMT exhibit differences in the methylation of protocatechuic acid under similar reaction conditions. While the former produced an equal mixture of meta and para methylated products^[30]^, StrAOMT was clearly selective towards the *meta* isomer vanillic acid. This suggests that structural elements distant from the active site may as well influence the regioselectivity of methylation. One such element might be the variable loop connecting the β5 strand with the α8 helix. Originally spotted as the main difference between animal and plant COMTs, this “insertion loop” is believed to provide a scaffold for the binding of caffeoyl-CoA in specialized plant enzymes^[31]^. A somewhat extended and highly variable loop is also found in the structures of all bacterial COMTs, including StrAOMT. However, its implications in the substrate specificity of bacterial enzymes remain unclear and probably do not involve the binding of CoA^[6]^. Curiously, DesAOMT is the first characterized bacterial COMT that possesses a short, animal COMT-like β5-α8 loop. Overall, regioselectivity of DesAOMT and StrAOMT appears to be dictated by a complex interplay between the chemical natural of the substrate, structural elements of the enzyme and reaction conditions, which agrees with reports for other COMTs^[12,14,23]^. The structures of StrAOMT and DesAOMT may be used for deeper investigation of the COMT reaction mechanism and protein engineering efforts to modulate regioselectivity or affinity towards selected substrates. The latter possibility is particularly intriguing with 5-hydroxyflavonoids and 2-hydroxybenzoic acids, which were discovered to be accepted by two bacterial catechol OMTs in this study.

## Conclusion

We performed thorough *in vitro* characterization of two promiscuous Class I O-methyltransferases from bacteria and determined their substrate and product scope, methylation regioselectivity and crystal structures. Both enzymes operated in a broad temperature and pH range, exhibited tolerance to organic solvents and methylated a broad range of natural product scaffolds with differing regioselectivies, which makes them excellent candidates for pathway engineering and combinatorial biosynthesis applications.

Our findings suggest that with optimized reaction conditions or further enzyme and pathway engineering, DesAOMT and StrAOMT could provide a sustainable alternative for the production of several natural products of demonstrated pharmaceutical relevance, such as (iso)fraxidin, iso(scopoletin), chrysoeriol, alizarin 1-methyl ether and 2-methoxyestradiol. All of these compounds are currently sourced either from producer plants or through chemical synthesis.

We found that StrAOMT and, to some extent, DesAOMT can methylate *o*-hydroxybenzoic acids and 5-OH flavonoids, which have not been associated with this class of enzymes before. To the best of our knowledge, this is the first report of the enzymatic synthesis of the 5-O-methyl ethers of naringenin, chrysin and pinocembrin. The latter, better known as alpinetin, is a rare flavonoid with demonstrated potential for the treatment of acute colitis^[32]^, among other conditions. As the native biosynthetic pathway for alpinetin remains unknown, and no flavonoid-5-OMTs have been described in the literature, StrAOMT is a good candidate for directed evolution efforts towards the improved enzymatic production of this and other 5-O-methylated flavonoids. The structural insights generated in this study may facilitate rational engineering of StrAOMT and DesAOMT towards the improved turnover of non-natural COMT substrates to valuable pharmaceuticals. Last but not least, the findings from our substrate scope analysis may provide inspiration for the development of new human COMT inhibitors.

## Experimental section

### Expression, purification and storage of DesAOMT and StrAOMT

The genes encoding DesAOMT and StrAOMT were subcloned into pET-21b(+) with a C-terminal 6xHis-tag. Plasmids harbouring the OMT genes were transformed into chemically competent *E. coli* BL21(DE3) and maintained on selective LB agar containing 100 mg/mL ampicillin. A starter culture was inoculated from a single colony (5 mL, LB with ampicillin) and incubated at 37°C (180 rpm, overnight). The main culture was inoculated from the starter culture (1:100) into auto-induction medium (2% w/v tryptone, 0.5% w/v yeast, 0.5% w/v sodium chloride, 25 mM disodium hydrogen phosphate dihydrate, 25 mM potassium dihydrogen phosphate, 0.6% v/v glycerol, 0.05% w/v glucose, 0.0128% w/v lactose) and incubated at 37°C (180 rpm, 2h), after which the temperature was lowered to 18°C (180 rpm, overnight). All following steps were performed with chilled buffers. The cells were harvested by centrifugation (15 min, 3428 x g) and the pellet was resuspended in 5 volumes of the lysis buffer (buffer A including one EDTA-free protease inhibitor tablet (Roche); buffer A: 50 mM Tris/HCl pH 7.5, 500 mM NaCl, 20 mM imidazole). The cell suspension was lysed by sonication (40% duty cycle, 6 cycles of 30 s ON/30 s OFF) and cleared by centrifugation for 60 min at 25000 x g. The supernatant was loaded onto a HisTrap HP Ni-NTA column (GE Healthcare, USA) connected to an ÄKTA Pure system (Amersham Bioscience, Uppsala, Sweden) and eluted with a linear gradient 0-100% of buffer B (50 mM Tris/HCl pH 7.5, 500 mM NaCl, 500 mM imidazole). Elution fractions corresponding to the protein peak were analysed by SDS PAGE. Fractions with low protein background were pooled and subjected to size-exclusion chromatography on a Superdex 75 pg column (GE Healthcare, USA) in the storage buffer (DesAOMT: 10 mM Tris/HCl pH 7.4, 20 mM NaCl, 0.2 mM MgCl_2_, 5 mM BME; StrAOMT: 10 mM HEPES pH 7, 200 mM NaCl, 0.2 mM MgCl_2_, 10 mM DTT, 5% v/v glycerol). The protein concentration was determined by absorbance at 280 nm (NanoDrop, ThermoFisher Scientific, USA) before the purified enzymes were aliquoted and flash-frozen with liquid nitrogen for storage at -80°C.

### Differential scanning fluorimetry (DSF)

For each screening, 0.5 mL of a 1–2 mg/mL protein sample was mixed with 2.5 µL of SYPRO® orange dye (ThermoFisher Scientific, USA) and aliquoted at 5 µL before being mixed with 45 µL of the respective screening buffer. Thermal stability of the protein samples was measured in a CFX96 Dx Real-Time qPCR instrument (Bio-Rad, Hercules, CA, USA); program: 20°C for 2 min, 20–95°C over 117 min. Protein melting temperatures (Tm) under the different buffer conditions were determined from the maximum value of the first derivative of the melting curve.

### Activity tests

The initial conditions for the *in vitro* OMT reaction were adapted from Siegrist et al.^[23]^ and included 50 mM HEPES/NaOH pH 7, 20 mM MgCl_2_, 1 mM SAM, 0.5 mM substrate (from 80 mM stock in DMSO)) and 5 µM enzyme in a total volume of 42 µL. The reactions were started by the addition of the enzyme (or water for the “no OMT” control). The reactions were incubated for 1h at 30°C unless stated otherwise, quenched with HClO_4_ (final 2% v/v), centrifuged and stored at 4°C until analysed.

To determine the optimal temperature for the OMT activity, the reactions were incubated in a temperature range of 30-60°C. All subsequent reactions were incubated at 37°C. To investigate metal dependence of the OMTs, different cations (Mg^2+^, Ca^2+^, Mn^2+^, Co^2+^, Ni^2+^, Zn^2+^, Cu^2+^) or EDTA were added to the reaction at a concentration of 2 mM alongside a no additive (“none”) control. To study the pH and buffer effects, the reactions were incubated with a 5-fold excess of SAM and 50 mM of the respective buffer (NaPi pH 6; HEPES pH 7; Tris-HCl pH 7.5, 8, 8.5; Gly-NaOH pH 9, 10, 11). 20 mM Tris-HCl (pH 7.5) was used for subsequent analyses. For solvent tolerance studies, 0–20% (v/v) of DMSO, methanol or acetonitrile was added to the reaction mix right before the addition of the enzyme. For substrate scope studies, the reactions were incubated for 16 h at 37°C with the addition of 10-20% (v/v) DMSO.

### Steady-state kinetics

Kinetic analyses were performed using 200 nM StrAOMT or 500 nM DesAOMT and 10 to 1000 µM substrate in a reaction mix consisting of 20 mM Tris-HCl pH 7.5, 20 mM MgCl_2_ and 1 mM SAM at 37°C. For the estimation of the initial velocity using linear regression, the reactions were quenched after 5, 10 and 15 min with HClO_4_ (final 2% v/v), centrifuged and stored at 4°C before HPLC analysis. The peak areas were integrated and converted to concentrations in µM based on calibration curves with the authentic standards. The apparent initial velocities of all experiments performed in triplicate were plotted against substrate concentrations using GraphPad Prism 8 and the apparent K_m_ and k_cat_ constants were determined by non-linear regression with the Michaelis-Menten equation. A full report of the regression statistics is given in Table S1.

### Analysis and quantification of OMT reaction products

The supernatants of the quenched OMT reactions were analysed by reversed-phase HPLC (instrument: Shimadzu LC-10AT; autosampler: HiP sampler G1367A, T = 4°C, 10 µL injection; flow rate: 1 mL/min; column: Agilent Zorbax Eclipse XDB-C18 80Å, 4.6 × 150 mm, 5 µm, T = 30°C; detector: SPD-20A photodiode array detector (PDA), λ = 275 nm (SAM/SAH; alizarin, DHICA, carnosic acid, pinocembrin, 2-hydroxyestradiol and their methylated products) and λ = 310 nm (all other substrates and their methylated products); solvents A: water with 0.1% TFA, solvent B: ACN with 0.1% TFA; gradient: 10–28% B over 12.5 min; 28–100% B over 9.5 min; 100– 10% B over 2 min; 10% B for 3 min.). For analysing the samples of the steady-state kinetics, a shorter program was used: 10–13% B over 2.5 min; 13–25% B over 1.5 min; 25–35% B over 2 min; 35-65% B over 2 min; 65–100% B over 1 min; 100–10% B over 3 min; 10% B for 3 min. Product peaks were identified by comparing the retention times to authentic standards (where available). The peak areas were integrated and converted to concentrations in µM based on calibration curves with the authentic standards (where available). Regioisomeric excess (RE) of the reaction was calculated using the formula: (RE=(c[meta]-c[para])/(c[meta]+c[para])*100).

The identity of reaction products was furthermore confirmed by HPLC-coupled mass spectrometry (LCMS) with a Waters Acquity Arc UHPLC-MS equipped with a 2998 PDA, and a QDa single quadrupole mass detector. The samples were separated over an XBridge BEH C18 3.5 µm 2.1 × 50 mm column with a concentration gradient (solvent A: water + 0.1% formic acid, and solvent B: acetonitrile + 0.1% formic acid) at a flow rate of 0.5mL/min (2 μL injections). The following gradient was used: 5% B for 2 min, 5–90% B over 3 min; 90% B for 2 min; 5% B for 3 min.

### Protein crystallization

The sitting-drop vapor diffusion method was applied for crystallization of DesAOMT and StrAOMT at 18°C. Sparse-matrix screening was carried out using the commercial kits JCSG plus, PACT premier, Morpheus, PGA and the MIDAS plus screen (Molecular Dimensions Ltd., UK). Reservoir solution and freshly prepared protein were mixed at a ratio of 1:1 μl. DesAOMT was used at a concentration of 8 mg/ml in 10 mM HEPES pH 7.0, 20 mM NaCl, 0.2 mM MgCl_2_, and 5 mM BME. After 5-6 days, a tetragonal bipyramid-shaped crystal appeared in the well containing reservoir solution of 0.1 M MES/Imidazole pH 6.5, 0.03 M MgCl_2_, 0.03 M CaCl_2_, 20% (v/v) glycerol and 10% (w/v) PEG4000. StrAOMT was concentrated to 13 mg/ml in 10 mM HEPES pH 7, 200 mM NaCl, 0.2 mM MgCl_2_, 10 mM DTT and 5% v/v glycerol. Monoclinic crystals of StrAOMT were obtained in a drop containing reservoir solution of 0.1 M MIB (Malonic acid, Imidazole, Boric acid) pH 6.0 and 25% (w/v) PEG1500.

For co-crystallization of StrAOMT with SAH, the protein was incubated with 2.67 mM SAM (from a 32 mM stock containing 5 mM H_2_SO_4_ and 10% (v/v) EtOH) and 2.5 mM caffeic acid (from a 1 M DMSO stock) for half an hour at room temperature prior to crystallization. Reservoir solution for the StrAOMT-SAH complex contained 0.2 M ammonium formate, 10% (w/v) polyvinylpyrrolidone, and 20% (w/v) PEG 4000. Crystals were harvested using nylon loops 3-4 days before the diffraction experiments. The crystals were briefly immersed in cryoprotectants made from reservoir solutions supplemented with glycerol (20-30% (v/v)) and flash-cooled in liquid nitrogen.

### Diffraction data collection, structure determination and refinement

Diffraction data were collected at beamline P11 at the PETRA III (DESY, Hamburg, Germany) at 100 K. Auto-processing of diffraction data was carried out with XDSAPP^[33]^. Aimless in the CCP4 suite^[34]^ was used for further truncation of the data and analysis of the merging statistics.

The crystal structures were solved through molecular replacement using the MOLREP program^[35]^. The apo StrAOMT model was solved using a search model of the monomer of O-methyltransferase from *Bacillus cereus* (PDB entry: 3DUW). Subsequently, the refined model of the apo-StrAOMT monomer served as a search model for the holo-structure of StrAOMT using the DIMPLE pipeline^[36]^. Finally, the AlphaFold model (Q1JXV1) was used to solve the DesAOMT structure. All structures were subjected to iterative cycles of refinement and model building using REFMAC5^[37]^ and Coot^[38]^. Automatically determined TLS parameters were used in REFMAC5 for DesAOMT on account of the significant anisotropy detected.

### PDB deposition

All three structures were deposited in the PDB under accession codes 8C9V (DesAOMT), 8C9T (apo-StrAOMT) and 8C9S (holo-StrAOMT).

### Bioinformatic analyses

Multiple sequence alignments were performed with mafft v7.505^[39]^ (parameters: --maxiterate 1000 – genafpair). Gene neighbourhoods of DesAOMT and StrAOMT and their sequence homologs were visualized with the EFI-GNT webtool^[22]^ using the Single Sequence Blast option and a neighbourhood window size of 20 genes.

## Acknowledgements

The authors acknowledge DESY (Hamburg, Germany) and EMBL Hamburg for the provision of experimental facilities at PETRA III. NS, SD and KH are grateful to Dr. Robbert Cool for support with protein purification and Serj Koshian for the initial genome neighbourhood analysis of the OMTs.

## Author contributions

NS and KH conceived the study; NS and SD expressed and purified the proteins and performed biochemical characterization; NS and KH analysed biochemical data; LZ and RO performed crystallization and diffraction experiments; LZ, RO and MG determined and refined the crystal structures; NS and KH wrote the manuscript with contributions from LZ, RO and MG; all authors have read and approved the final version of the manuscript.

