## Supplementary Figures and Tables for "Structural characterization and extended substrate scope analysis of two Mg^2+^-dependent O-methyltransferases from bacteria"

### Supporting tables

Table S1. Michaelis-Menten kinetic parameters of DesAOMT and StrAOMT

| **Enzyme** | **DesAOMT** | **StrAOMT** |
|---|---|---|
| Best-fit values | | |
| Et, µM | = 0,5000 | = 0,2000 |
| $k_{cat}$, $min^{-1}$ | 0,9022 | 2,308 |
| $K_m$, µM | 63,85 | 122,5 |
| $V_{max}$, uM/min | = 0,4511 | = 0,4617 |
| **95% CI (profile likelihood)** | | |
| $k_{cat}$, $min^{-1}$ | 0,7901 to 1,037 | 2,144 to 2,495 |
| $K_m$, µM | 43,35 to 94,63 | 95,92 to 158,4 |
| **Goodness of Fit** | | |
| Degrees of Freedom | 16 | 15 |
| R squared | 0,9257 | 0,9814 |
| Sum of Squares | 0,02367 | 0,006269 |
| Sy.x | 0,03847 | 0,02044 |

Table S2. Diffraction data collection, structure determination, and refinement statistics. Values in brackets refer to the highest resolution shell.

| **Data set** | | **DesAOMT** | **StrAOMT** | **StrAOMT+SAH** |
|---|---|---|---|---|
| PDB entry | | 8C9V | 8C9T | 8C9S |
| Diffraction source | | beamline P11, DESY | | |
| Wavelength (Å) | | 1.0332 | | |
| Temperature (K) | | 100 | | |
| Detector | | EIGER2 2X 16M | | |
| Rotation range per image (°) | | 0.1 | | |
| Total rotation range (°) | | 360 | | |
| Crystal-to-detector distance (mm) | | 200.483 | 200.201 | 200.643 |
| Space group | | P43 21 2 | P1 21 1 | P1 21 1 |
| Unit cell parameters | a, b, c (Å) | a = 44.791 | a = 46.457 | a = 46.398 |
| | | b = 44.791 | b = 168.744 | b = 41.596 |
| | | c = 204.919 | c = 57.242 | c = 104.207 |
| | α, β, γ (°) | α = 90.00 | α = 90.00 | α = 90.00 |
| | | β = 90.00 | β = 101.487 | β = 96.564 |
| | | γ = 90.00 | γ = 90.00 | γ = 90.00 |
| Resolution, Å | | 44.79-1.50(1.53-1.50) | 46.71-1.50(1.53-1.50) | 46.09-1.8(1.84-1.80) |
| Number of observations | | 386106(13400) | 937293(32598) | 279208(15953) |
| Completeness, % | | 99.3(100.0) | 96.2(79.0) | 99.2(99.1) |
| Multiplicity | | 11.1(8.0) | 7.1(6.2) | 7.6(7.4) |
| Rmerge | | 0.043(1.447) | 0.051(0.278) | 0.158(1.406) |
| Average I/σ,I | | 23.0(1.2) | 21.6(5.7) | 9.2(1.5) |
| CC(1/2) | | 1.000(0.828) | 0.999(0.967) | 0.998(0.603) |
| Wilson B-factor, Å2 | | 21.5 | 12.9 | 18.5 |
| **Refinement statistics** | | | | |
| No. reflections, all/free | | 34687/1706 | 131918/6583 | 36669(1855) |
| Rfactor | | 0.171 | 0.167 | 0.162 |
| Rfree | | 0.217 | 0.194 | 0.211 |
| No. protein atoms | | 1449 | 6833 | 3398 |
| No. ligand atoms | | - | - | 64 |
| No. water atoms | | 99 | 712 | 173 |
| **Average B-factors, Å2** | | | | |
| Protein | | 34.5 | 14.9 | 23.67 |
| Ligands | | - | - | 20.25 |
| Water | | 39.74 | 24.55 | 29.39 |
| **RMSD from ideal values** | | | | |
| Bond lengths, Å | | 0.0144 | 0.0137 | 0.0096 |
| Bond angles, ° | | 1.773 | 1.825 | 1.528 |
| **Ramachandran plot** | | | | |
| Favoured, % | | 97.27% | 96.92% | 97.06% |
| Allowed, % | | 2.19% | 2.51% | 1.81% |
| Disallowed, % | | 0.54% | 0.57% | 1.13% |

Table S3: Structural alignment of DesAOMT and StrAOMT with known OMTs from PDB with the Dali server

| # | PDB ID and chain | Z | rmsd | lali | nres | %id | Ligands | Protein name | Donor organism |
|---|---|---|---|---|---|---|---|---|---|
| **StrAOMT** | | | | | | | | | |
| 1 | 3duw-A | 36.6 | 1 | 220 | 221 | 56 | SAH | BcOMT2 | *Bacillus cereus* ATCC 10987 |
| 2 | 7cvx-B | 36.2 | 1.2 | 221 | 221 | 52 | SAH, $Mg^{2+}$, 3,4-DHB | NkCOMT | *Niastella koreensis* GR20-10 |
| 3 | 5n5d-B | 35.8 | 1.3 | 224 | 224 | 51 | SAM | TomG | *Streptomyces regensis* |
| 4 | 3tfw-A | 35.6 | 1 | 220 | 221 | 53 | - | A6T8J6 | *Klebsiella pneumoniae* subsp. *pneumoniae* MGH 78578 |
| 5 | 6jcl-H | 34.6 | 1.5 | 216 | 217 | 56 | SAH, $Sr^{2+}$ | Rv0187 | *Mycobacterium tuberculosis* H37Rv |
| 6 | 6jcm-C | 32.5 | 1.6 | 214 | 215 | 57 | - | Rv0187 | *Mycobacterium tuberculosis* H37Rv |
| 7 | 3cbg-A | 27.1 | 2 | 209 | 219 | 31 | SAH, $Mg^{2+}$, ferulic acid, isoferulic acid | SynOMT | *Synechocystis* sp. PCC 6803 |
| **DesAOMT** | | | | | | | | | |
| 1 | 2hnk-C | 26.1 | 1.8 | 181 | 232 | 31 | SAH | LiOMT | *Leptospira interrogans* |
| 2 | 3tr6-A | 25.2 | 1.8 | 178 | 223 | 30 | SAH, $Ni^{2+}$ | Q83D22 | *Coxiella burnetii* |
| 3 | 5x7f-A | 25.1 | 2 | 177 | 199 | 31 | SAM | Rv1220c | *Mycobacterium tuberculosis* H37Rv |
| 4 | 5log-A | 25.1 | 1.9 | 180 | 223 | 32 | SAH, $Mg^{2+}$, L-dopamine | SafC | *Myxococcus xanthus* |
| 5 | 6jcl-F | 24.3 | 1.8 | 174 | 217 | 28 | SAH, $Sr^{2+}$ | Rv0187 | *Mycobacterium tuberculosis* H37Rv |
| 6 | 1sui-A | 24.3 | 1.9 | 179 | 228 | 28 | SAH, $Ca^{2+}$, feruloyl-CoA | Q40313 | *Medicago sativa* |
| 7 | 3c3y-B | 24.2 | 2 | 179 | 226 | 25 | $Ca^{2+}$, SAH | PFOMT | *Mesembryanthemum crystallinum* |

### Supporting figures

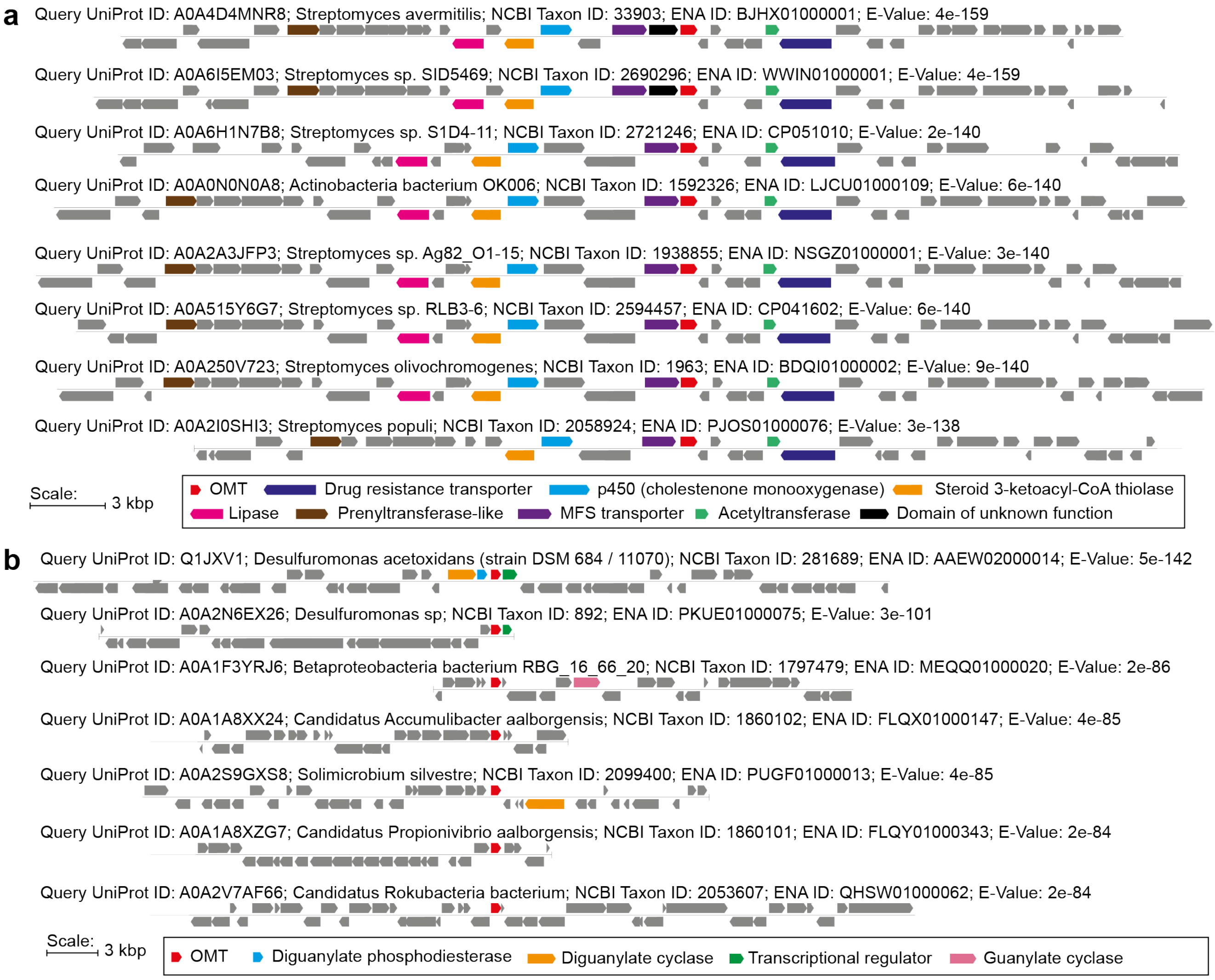

Figure S1. Gene neighbourhood diagrams of the OMTs: a) DesAOMT and sequence homologs; b) StrAOMT and sequence homologs. Selected putative functionalities are colour-coded.

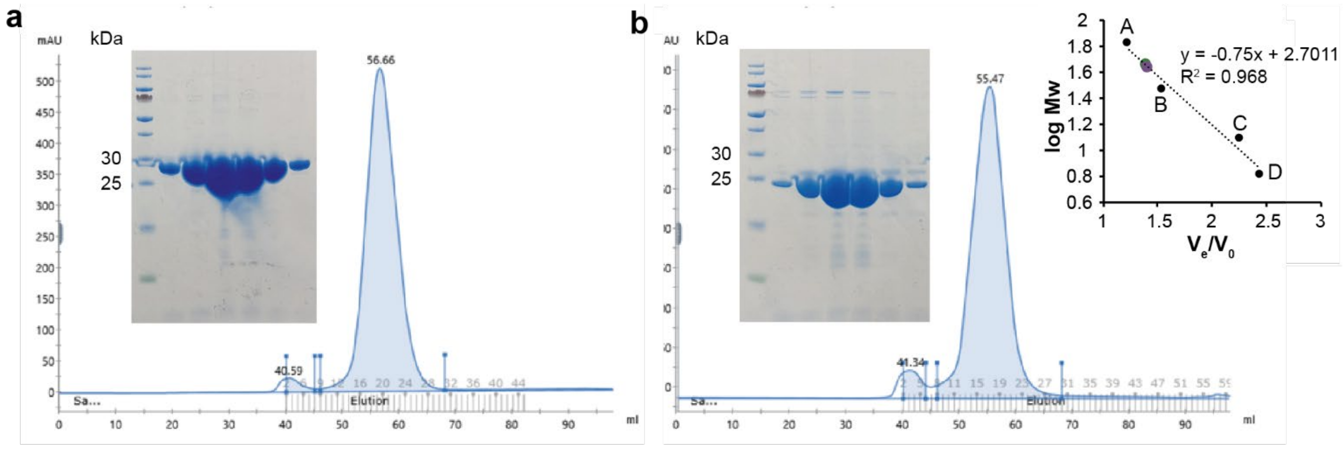

Figure S2. Purification and size-exclusion chromatography analysis of a) StrAOMT and b) DesAOMT. Column calibration was performed in DesAOMT storage buffer with the following protein standards: A – bovine serum albumin (66 kDa), B – carbonic anhydrase (29 kDa), cytochrome C (12.4 kDa), aprotinin (6.5 kDa). The purple and green dots on the calibration represent StrAOMT and DesAOMT, respectively.

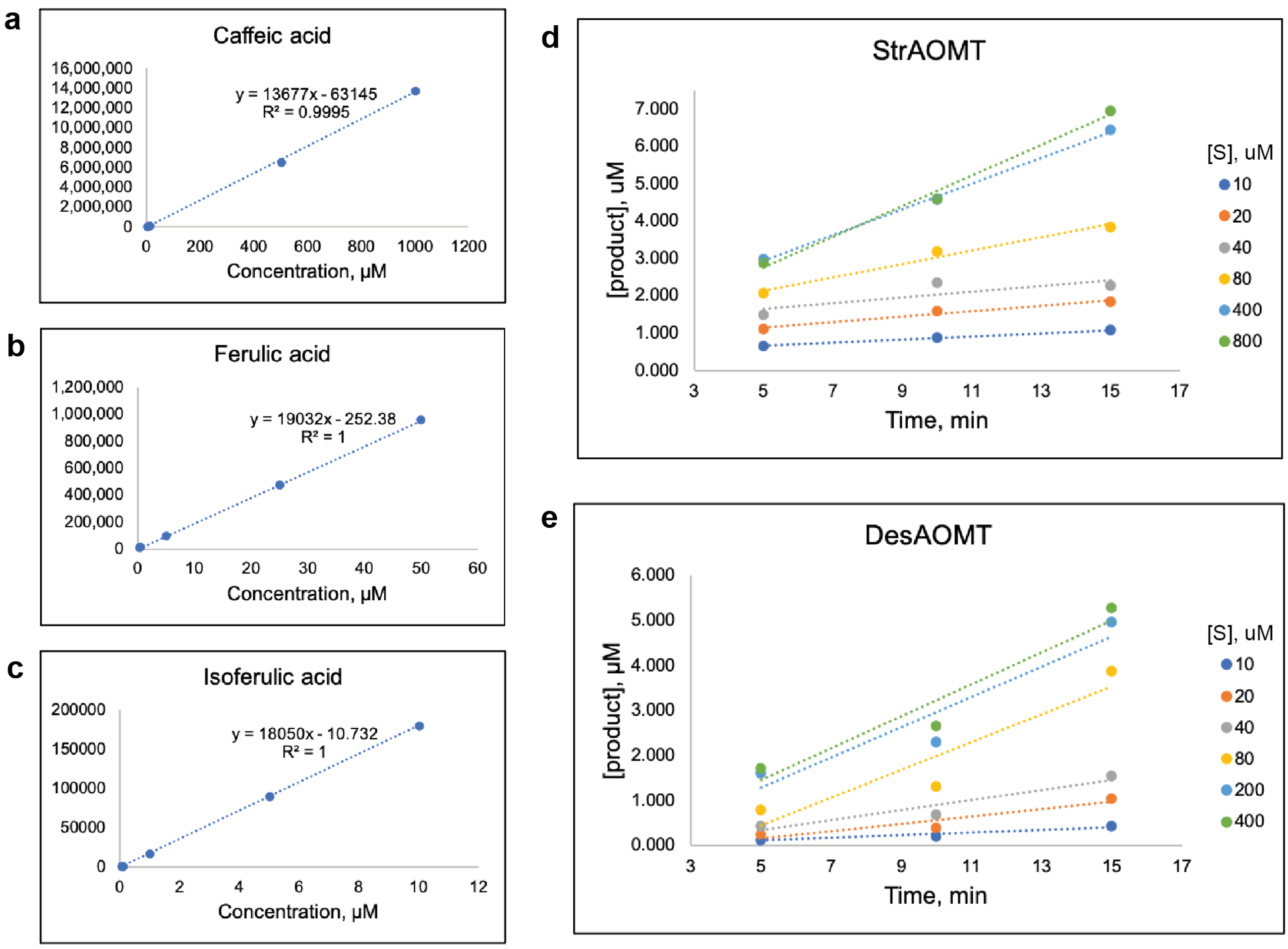

Figure S3. Calibration plots of a) caffeic acid, b) ferulic acid and c) isoferulic acid dissolved in the reaction buffer (20 mM Tris-HCl pH 7.5, 20 mM $MgCl_2$) and analysed by HPLC; estimation of the initial velocity of d) StrAOMT and e) DesAOMT at different substrate concentrations. Data are represented as means. The compounds were detected at 310 nm and the analysis was performed as described in the method section.

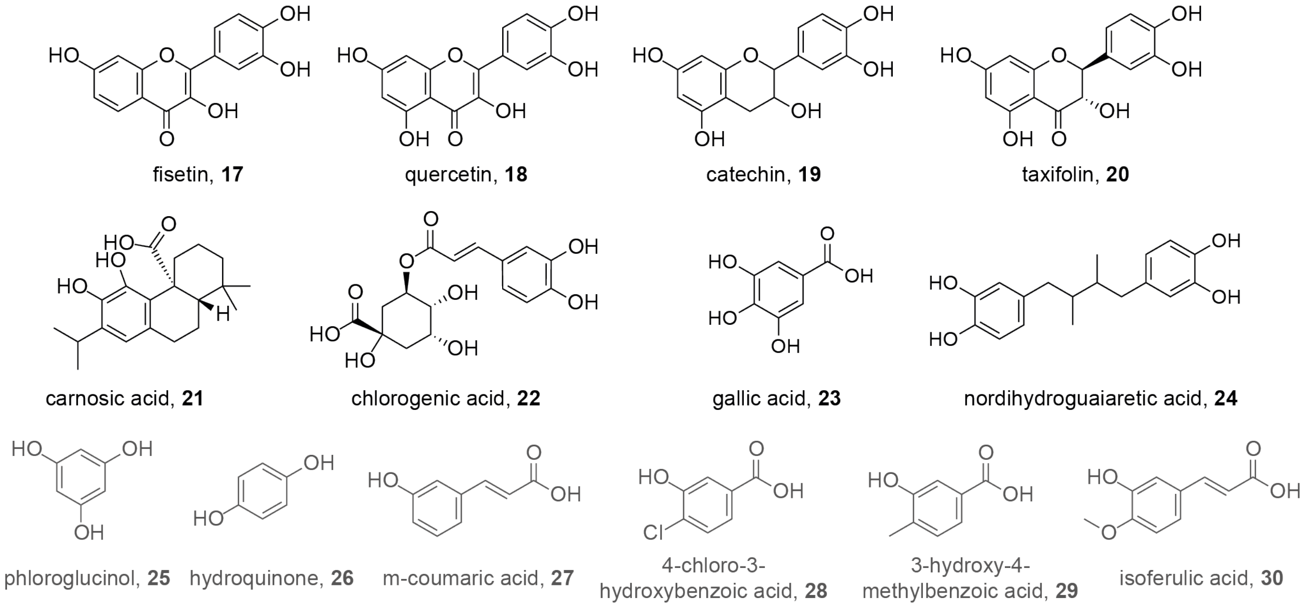

Figure S4. Extended substrate panel. Substrates that were accepted (**17-24**) are depicted in black; substrate that were not accepted (**25-30**) are depicted in grey.

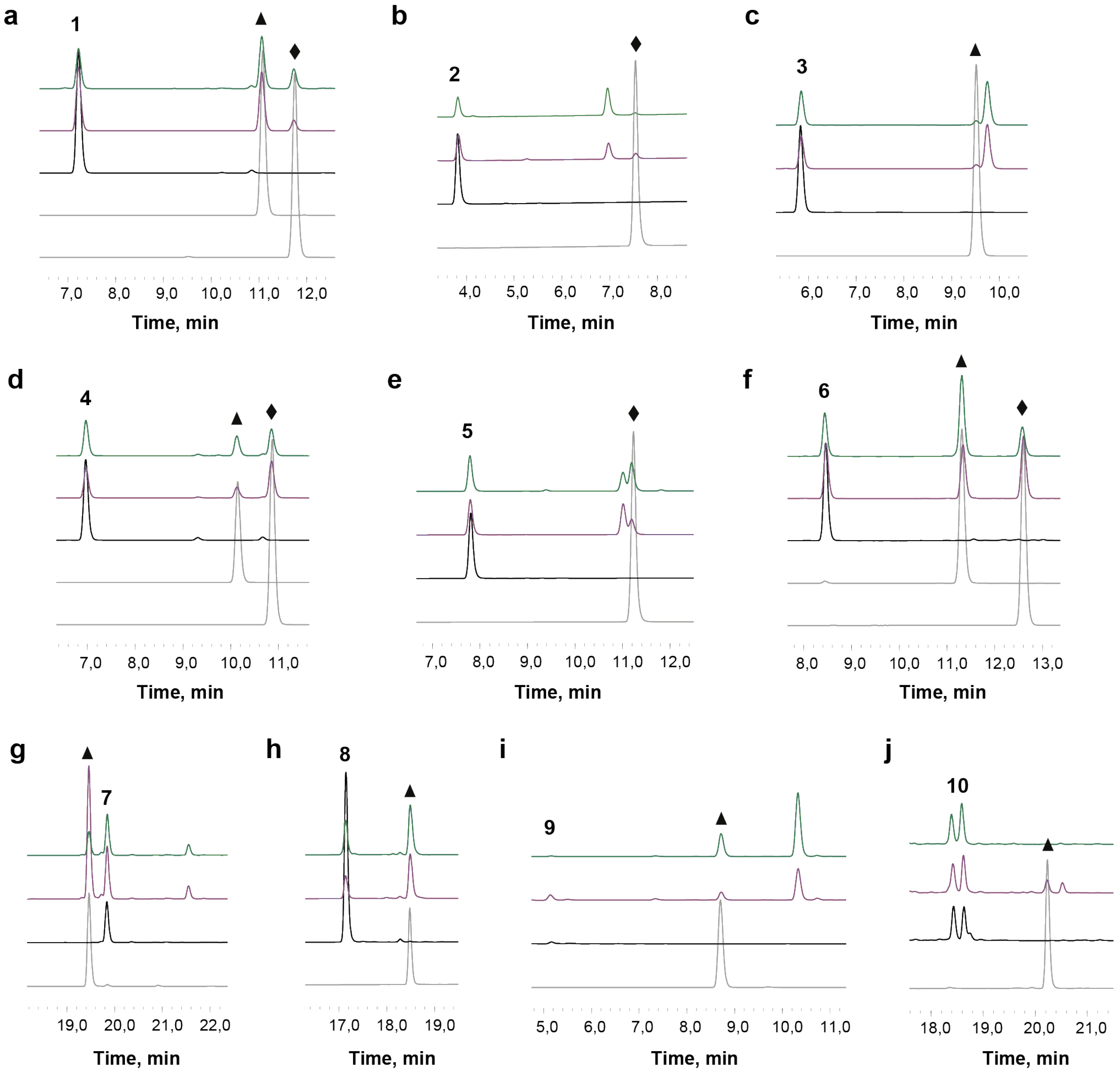

Figure S5. HPLC traces of the methylation reactions with catechol-like substrates **1-10** catalysed by DesAOMT and StrAOMT. Blue lines – DesAOMT reaction, red lines – StrAOMT reaction, black lines – "no enzyme" control, grey lines – authentic standards of the potential products: a) substrate **1**, standards: ferulic acid (triangle), isoferulic acid (diamond); b) substrate **2**, standard: isovanillic acid (diamond); c) substrate **3**, standard: isovanillin (triangle); d) substrate **4**, standards: isoscopoletin (triangle), scopoletin (diamond); e) substrate **5**, standard: 7-methoxy-8-hydroxycoumarin; f) substrate **6**, standards: isofraxidin (triangle), fraxidin (diamond); g) substrate **7**; standard: alizarin 2-methyl ether (triangle); h) substrate **8**, standards: chrysoeriol (triangle); i) substrate **9**, standard: 5-hydroxy-6-methoxyindole-carboxylic acid (triangle); j) substrate **10**, standard: 2-methoxyestradiol (triangle).

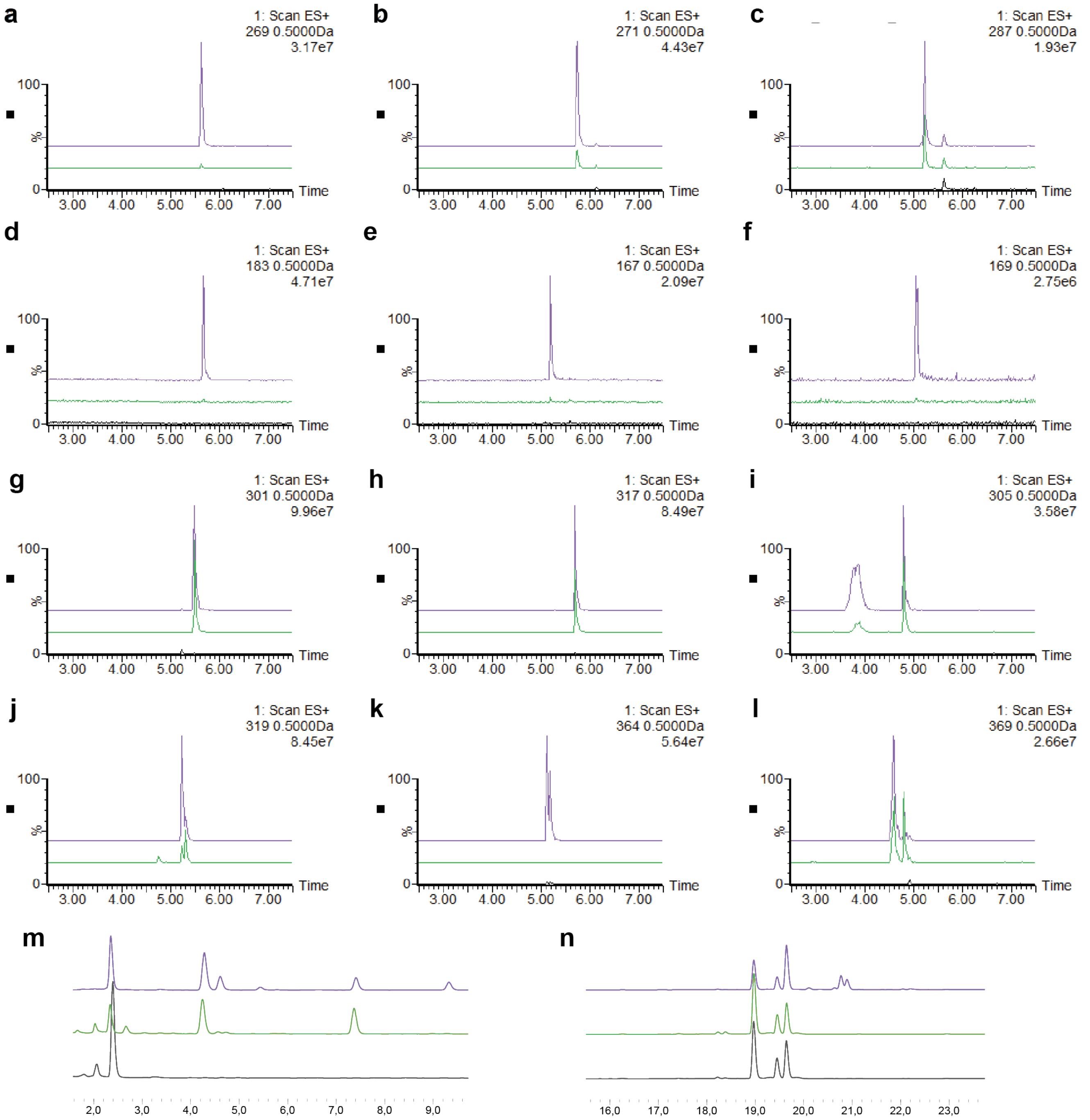

Figure S6. Methylation reactions with substrates **11**-**24** catalysed by DesAOMT (green) and StrAOMT (purple). Extracted ion chromatograms of the reaction products with a) substrate **11**; b) substrate **12**; c) substrate **13**; d) substrate **14**; e) substrate **15**; f) substrate **16**; g) substrate **17**; h) substrate **18**; I) substrate **19**; j) substrate **20**; k) substrate **21**; l) substrate **22**; HPLC traces of the methylation reactions with m) substrate **23**; n) substrate **24**. The following mass-to-charge ratios (m/z) were detected in positive ion mode: mono-methylated **11**, m/z = 269.0 (theoretical m/z = 269.0813, calculated for $C_{16}H_{13}O_4$), mono-methylated **12**, m/z = 271.1 (theoretical m/z = 271.0970, calculated for $C_{16}H_{15}O_4$); mono-methylated **13**, m/z = 286.9 (theoretical m/z = 287.0919, calculated for $C_{16}H_{15}O_5$); mono-methylated **14**, m/z = 183.1 (theoretical m/z = 183.0657, calculated for $C_9H_{11}O_4$); mono-methylated **15**, m/z = 167.1 (theoretical m/z = 167.0708, calculated for $C_9H_{11}O_3$); mono-methylated **16**, m/z = 169.0 (theoretical m/z = 169.0500, calculated for $C_8H_9O_4$); mono-methylated **17,** m/z = 300.9 (theoretical m/z = 301.0712, calculated for $C_{16}H_{13}O_6$); mono-methylated **18,** m/z = 317.0 (theoretical m/z = 317.0661, calculated for $C_{16}H_{13}O_7$); mono-methylated **19,** m/z = 304.8 (theoretical m/z = 305.1025, calculated for $C_{16}H_{17}O_6$); mono-methylated **20,** m/z = 318.9 (theoretical m/z = 319.0817, calculated for $C_{16}H_{15}O_7$); **21,** m/z = 364.2 (theoretical m/z = 347.2222, calculated for $C_{21}H_{31}O_4$); mono-methylated **22,** m/z = 369.17 (theoretical m/z = 369.1185, calculated for $C_{17}H_{21}O_9$).

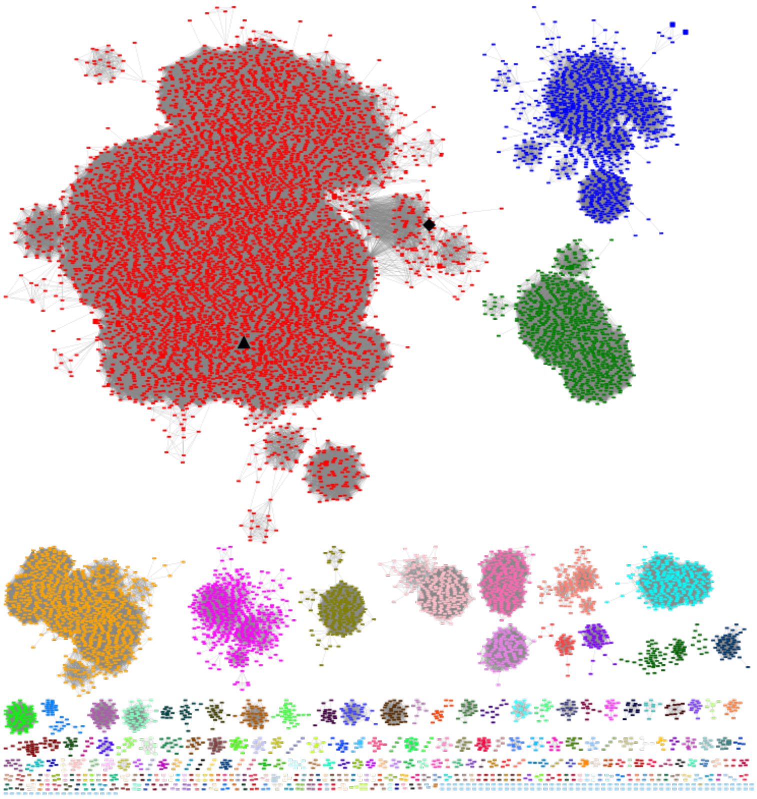

Figure S7. Sequence similarity network of the OMTs at 45% sequence identity cut-off. StrAOMT is marked with a triangle, DesAOMT is marked with a diamond.

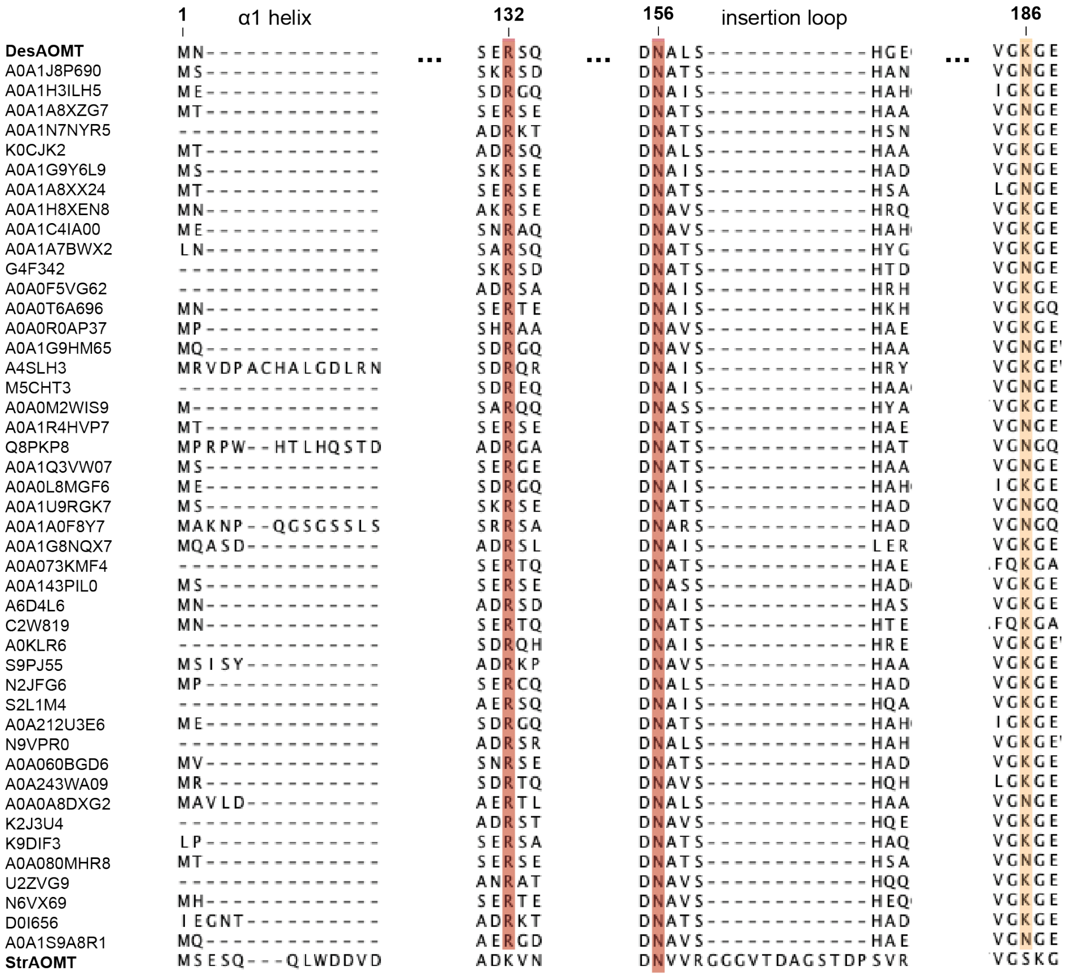

Figure S8. Alignment of the sequences from the DesAOMT subcluster of the OMT sequence similarity network (55% sequence identity cut-off; residue numbering from DesAOMT). The sequences are named after their UniProt IDs. StrAOMT is included for the comparison of the α1 helix and the insertion loop regions.

```
Q82B68|StrAOMT      --------------------------------------------MSESQQ--LWDDVDDY    14
Q739U3|BcOMT2       ---------------------------------------------MSMIE--TWTAVDQY    13
G8T6H8|NkCOMT       ----------------------------------------------MNNQ--IFESVDHY    12
C0LTS8|TomG         ----------------------------------------------MTQH--QWSAVDGY    12
O07431|Rv0187       -------------------------------------------MGMDQQP--NPPDVDAF    15
Q55813|SynOMT       -------------------------------------------MGKGITG--FDPSLYSY    15
Q1JXV1|DesAOMT      ----------------------------------------------MNK------ELHQL    8
Q8F8Y3|LiOMT        -------------------------------------------MSRKNIS--LTESLEEY    15
Q50859|SafC         --------------------------------------------MIHHVE--LTQSVLQY    14
Q6YI95|PFOMT        -------------------------------MDF------AVMKQVKNTGLLQSEELCQY    23
P22734|RnCOMT       MPLAAVSLGLLLLALLLLLRHLGWGLVTIFWFEYVLQPVHNLIMGDTK-----EQRILRY    55
                                                                            :

Q82B68|StrAOMT      FTTLLA--PEDEALTAALRDSDAAGLPHINVAPNQGKLLQLLAEIQGARRILEIGTLGGY    72
Q739U3|BcOMT2       VSDVLI--PKDSTLEEVLQVNAAANLPAHDVSPTQGKFLQLLVQIQGARNILEIGTLGGY    71
G8T6H8|NkCOMT       ISDLLG--YEDDALLAATNSLAEAGMPAISVSPNQGKFLQLLAQLCQAKNILELGTLAGY    70
C0LTS8|TomG         FDTVLG--DEDPALAAAAQAHIGFDLPDIGVSRSQGKLLHLLALVQRAERILEIGTFGGY    70
O07431|Rv0187       LDSTLV--GDDPALAAALAASDAAELPRIAVSAQQGKFLCLLAGAIQARRVLEIGTLGGF    73
Q55813|SynOMT       LQSISA--DDSFYLAQLRRETAHLPGAPMQISPEQAQFLGLLISLTGAKQVLEIGVFRGY    73
Q1JXV1|DesAOMT      LCELE------EFGQANDEKTEDRAQRMLNITPDTGEFLAVLVRAMNARRVLEIGTSNGY    62
Q8F8Y3|LiOMT        IFRNSV--REPDSFLKLRKETGTLAQANMQISPEEGQFLNILTKISGAKRIIEIGTFTGY    73
Q50859|SafC         IRDSSV--RDNDILRDLREETSKLPLRTMQIPPEQGQLLSLLVRLIGARKTLEVGVFTGY    72
Q6YI95|PFOMT        ILRTSVYPREAGFLKELREANESHPDSYMSTSPLAGQLMSFVLKLVNAKKTIEVGVFTGY    83
P22734|RnCOMT       VQQNA-KPGDPQSVLEAIDTYCTQKEWAMNVGDAKGQIMDAVIREYSPSLVLELGAYCGY    114
                    .                                  .:::  :         :*:*.  *:

Q82B68|StrAOMT      STIWLGRALP-RDGRLISFEYDAKHAEVARRNLARAGLDGISEVRVGPALESLPKLADER    131
Q739U3|BcOMT2       STIWLARGLS-SGGRVVTLEASEKHADIARSNIERANLNDRVEVRTGLALDSLQQIENEK    130
G8T6H8|NkCOMT       STIWMARALP-KNGRLITLEYDPKHAAVAQKNIDRAGLTSQVQIRTGKAIDILPQLVEEG    129
C0LTS8|TomG         STIWLARALP-PGGRLVTIEWERSFAETAAGHLARAGVADRVDQRVGRALDILPDLAKTG    129
O07431|Rv0187       STIWLARGAG-PQGRVVTLEYQPKHAEVARVNLQRAGVADRVEVVVGPALDTLPTLAG--    130
Q55813|SynOMT       SALAMALQLP-PDGQIIACDQDPNATAIAKKYWQKAGVAEKISLRLGPALATLEQLTQGK    132
Q1JXV1|DesAOMT      STLWLADAVSAIDGSVTTVEYAEQKYRLAQKNFSRTSLAHRIDAILGDAGTILGNADD--    120
Q8F8Y3|LiOMT        SSLCFASALP-EDGKILCCDVSEEWTNVARKYWKENGLENKIFLKLGSALETLQVLIDSK    132
Q50859|SafC         STLCAALALP-ADGRVIACDLSEEWVSIARRYWQRAGVADRIEVRLGDAHHSLEALVGSE    131
Q6YI95|PFOMT        SLLLTALSIP-DDGKITAIDFDREAYEIGLPFIRKAGVEHKINFIESDAMLALDNLLQGQ    142
P22734|RnCOMT       SAVRMARLLQ-PGARLLTMEMNPDYAAITQQMLNFAGLQDKVTILNGASQDLIPQLKKKY    173
                    * :  .       . :   :   .            .:        . :   :
                                         ♦                           ♦♦
Q82B68|StrAOMT      PEP------------FDLVFIDADKVNNPHYV--EWALKLTRPGSLIVVDNVVRGGGVTD    177
Q739U3|BcOMT2       YEP------------FDFIFIDADKQNNPAYF--EWALKLSRPGTVIIGDNVVREGEVID    176
G8T6H8|NkCOMT       AGP------------FDMIFIDADKPPYTEYF--QWALRLSRPGTLIVADNVIRDGKVLD    175
C0LTS8|TomG         EQP------------FDLVFIDANKPDTPEYF--DWALELSRPGGVIVVDNVVLGGAVAD    175
O07431|Rv0187       -GP------------FDLVFIDADKENNVAYI--QWAIRLARRGAVIVVDNVIRGGGILA    175
Q55813|SynOMT       PLPE-----------FDLIFIDADKRNYPRYY--EIGLNLLRRGGLMVIDNVLWHGKVTE    179
Q1JXV1|DesAOMT      -A------------VYDLIFLDSERSQYPGWW--PDLKRLLRPGGLLVVDNALSHGEQMA    165
Q8F8Y3|LiOMT        SAPSWASDFAFGPSSIDLFFLDADKENYPNYY--PLILKLLKPGGLLIADNVLWDGSVAD    190
Q50859|SafC         HRGT-----------FDLAFIDADKESYDFYY--EHALRLVRPGGLIILDNTLWSGKVAD    178
Q6YI95|PFOMT        ESE----------GSYDFGFVDADKPNYIKYH--ERLMKLVKVGGIVAYDNTLWGGTVAQ    190
P22734|RnCOMT       DVD-----------TLDMVFLDHWKDRYLPDTLLLEKCGLLRKGTVLLADNVIVPGTPDF    222
                                    *: *:*  :              * : * ::  **.:  *

Q82B68|StrAOMT      AGS-----TDPSVRGTRSALELIAEHPKLSGTAVQTVGSKGYDGFALARVLP--------    224
Q739U3|BcOMT2       NTS-----NDPRVQGIRRFYELIAAEPRVSATALQTVGSKGYDGFIMAVVKE--------    223
G8T6H8|NkCOMT       ENS-----TEPAVQGARRFNAMLGANTAVDATILQMVGVKEYDGMALAIVK---------    221
C0LTS8|TomG         SDD-----PDGGVRGMRRFHEKVAAEPRVSATTIQTVGAKGYDGFTLAVILPAPDDRPEA    230
O07431|Rv0187       ESD------DADAVAARRTLQMMGEHPGLDATAIQTVGRKGWDGFALALVR---------    220
Q55813|SynOMT       VDP-----QEAQTQVLQQFNRDLAQDERVRISVIPL-----GDGMTLALKK---------    220
Q1JXV1|DesAOMT      PFK-----------------ALVEADVEFTTCVVPV-----GKGEFLATRSA--------    195
Q8F8Y3|LiOMT        LSH-----QEPSTVGIRKFNELVYNDSLVDVSLVPI-----ADGVSLVRKR---------    231
Q50859|SafC         PSVV----GDPETDSLRRINAKLLTDERVDLSMLPI-----ADGLTLARKR---------    220
Q6YI95|PFOMT        PESEVPDFMKENREAVIELNKLLAADPRIEIVHLPL-----GDGITFCRRLY--------    237
P22734|RnCOMT       LAY---------VRGSSSFECTHYSS---YLEYMKV-----VDGLEKAIYQG--PSSPDK    263
                                                     :        .*
```

Figure S9. The amino acid sequences of StrAOMT and DesAOMT aligned with their structural homologs and sequences of other representative Class I COMTs including BcOMT2 from *Bacillus cereus*, NkCOMT from *Niastella koreensis*, TomG from *Streptomyces regensis*, Rv0187 from *Mycobacterium tuberculosis*, SynOMT from *Synechocystis* sp. Strain PCC 6803, LiOMT from *Leptospira interrogans*, OMT from *Coxiella burnettii*, SafC from *Myxococcus xanthus*, PFOMT, and RnCOMT (P22734) from *Rattus norvegicus*. Black diamonds mark metal binding residues. Conserved active site lysines are highlighted in pink; Asn and Asp of the catalytic triad are highlighted in blue. Substrate-binding residues are highlighted in grey.
